# Spatio-temporal interaction of immune and renal cells determines glomerular crescent formation in autoimmune kidney disease

**DOI:** 10.1101/2024.12.18.629206

**Authors:** Zeba Sultana, Robin Khatri, Behnam Yousefi, Nikhat Shaikh, Saskia L. Jauch-Speer, Darius P. Schaub, Jonas Engesser, Malte Hellmig, Arthur L. Hube, Varshi Sivayoganathan, Alina Borchers, Anett Peters, Anna Kaffke, Hans-Joachim Paust, Thiago Goldbeck-Strieder, Ulrich O. Wenzel, Victor Puelles, Elion Hoxha, Thorsten Wiech, Tobias B. Huber, Ulf Panzer, Stefan Bonn, Christian F. Krebs

**Author notes:** equal contribution (Zeba Sultana, Robin Khatri, Behnam Yousefi, Ulf Panzer, Stefan Bonn, Christian F. Krebs).

## Abstract

Rapidly progressive glomerulonephritis (RPGN) is the most aggressive group of autoimmune kidney disease with the worst prognosis. Anti-neutrophil cytoplasmic antibody (ANCA) associated vasculitis, anti-glomerular basement membrane (anti-GBM) and lupus nephritis are the most common causes of RPGN and are characterized by the formation of glomerular crescents and infiltration of leukocytes that eventually lead to glomerulosclerosis and kidney failure.

In this work, we used high-resolution spatial transcriptomics of 32 ANCA, 19 lupus nephritis, 6 anti-GBM, and 6 control patients to understand how intercellular signaling between immune and renal tissue cells leads to renal inflammation and glomerular injury. Using 3,218,210 immune and kidney cells, we observed that the biological pathways involved in the sequence of glomerular crescent formation are similar across the diseases. While innate immune cells infiltrated the glomerular compartment relatively early, later increases in adaptive immune cells were largely restricted to the periglomerular regions. These changes in immune cells temporally correlated with increases in glomerular parietal epithelial (PEC) and fibrotic mesangial cells, suggesting disease-relevant functional signaling between these immune and renal cells. Cell communication analysis revealed early disease PDGF signaling from epithelial and mesangial cells to PECs, causing their activation and proliferation. At later stages, TGF-β signaling from macrophages, T cells, epithelial cells, and mesangial cells to PECs triggered the expression of extracellular matrix components resulting in glomerulosclerosis.

Our results highlight a spatio-temporally conserved progression into glomerular crescents and sclerosis for ANCA, lupus nephritis, and anti-GBM disease, which is driven by consecutive PDGF and TGF-β signaling to PECs.

## Introduction

Rapidly progressive glomerulonephritis (RPGN) is a clinical syndrome characterized by a fast decline in kidney function accompanied by distinct histological features such as necrotizing glomerulonephritis and glomerular crescent formation. As a disease category, glomerulonephritis is among the most common causes for end-stage renal disease associated with a high morbidity and mortality for the individual patient and a high economic burden for our society. RPGN can develop in patients with systemic lupus erythematosus (SLE), ANCA-associated vasculitis (ANCA-GN), and anti-glomerular basement membrane antibody disease (anti-GBM). Glomerular crescents are typically formed by infiltrating immune cells and by parietal epithelial cells (PECs) that usually build the Bowman’s capsule and proliferate in crescentic glomerulonephritis (cGN). However, a comprehensive definition of infiltrating immune cells, their precise localization and the cellular signaling that result in PEC proliferation and ultimately in glomerulosclerosis remains missing.

To facilitate the spatial understanding of cellular and molecular disease processes, recent technical developments allow for the spatially-resolved, multiplex transcriptional analysis of tissues^1^. These available technologies such as MERSCOPE, CosMx, and Xenium measure mRNA using slightly different approaches including signal amplification processes^2, 3^. The Xenium platform relies on predefined oligonucleotide padlock probes that are circularized and amplified after binding to the matching mRNA in a PCR reaction, finally resulting in an augmented detection signal. This enables the measurement of gene transcription at high-resolution, which is important for the precise identification and spatial localization of specific cell types.

Here we use high-resolution spatial transcriptomics combined with multiplex spatial protein detection to investigate the development of glomerular crescents in the human kidney. We defined the glomerular niche, uncovered the cellular composition of the glomerulus as well as the adjacent area, and found PECs to be major mechanistic hubs in RPGN etiology. A comparative analysis of cell interactions in cGN versus control conditions revealed strong signaling from PECs to other cell types. Consequently, we investigated the signaling pathways targeting PECs that might contribute to their activation. The analysis identified upregulated PDGF signaling originating from various glomerular cells and TGF-β signaling from fibrotic mesangial and immune cells to be major activating signals for PECs. In combination with pseudo-time analysis, this defined the sequential upregulation of first PDGF and later TGF-β and the respective downstream signaling in PECs, resulting in cell proliferation and crescent formation, respectively. The identification of PDGF and TGF-β in the sequence of crescent formation might provide a novel basis for targeted therapies in glomerulonephritis.

## Results

### Patient selection and study design

To understand the molecular pathogenesis of RPGN, we conducted a comprehensive analysis of kidney samples from patients with lupus nephritis (SLE, n=19), ANCA-GN (n=32), and anti-GBM disease (n=6) and from healthy kidney tissue taken from tumor nephrectomies (Fig. 1A). All patients with RPGN were included in the Hamburg Glomerulonephritis Registry (Table 1). Gene expression profiling of 480 genes at high-resolution was performed on tissue sections using an *in situ* multiplexed mRNA assay (10x Xenium)^2, 4^. Kidney biopsy samples were distributed across 8 Xenium slides (Fig. 1A). Samples from different diseases were distributed across the slides to minimize potential systematic variations (Extended Data Fig. 1A).

**Figure 1:**
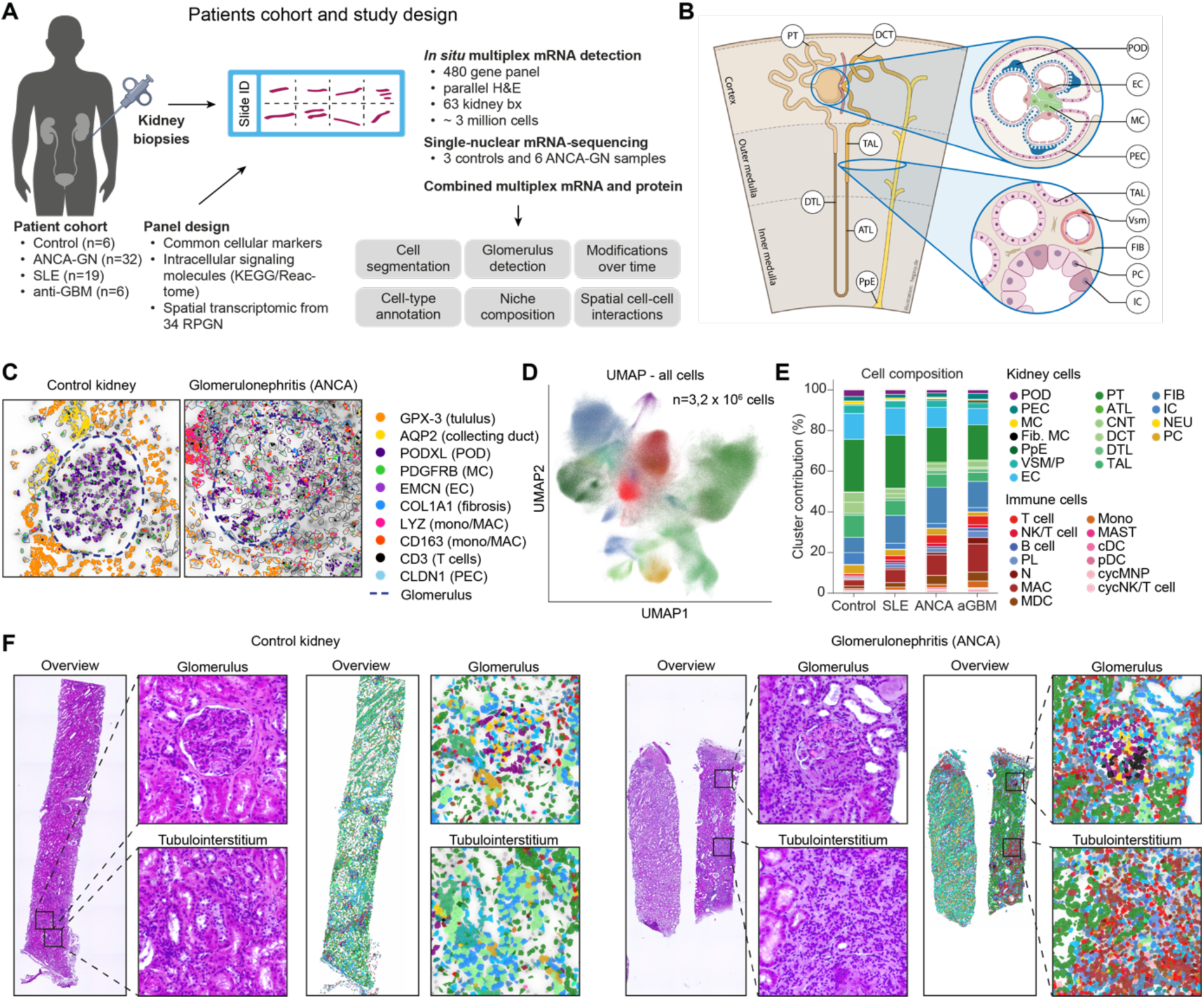
Major cell types in the human renal cortex in health and inflammation. **A,** Overview of the patient cohort from the Hamburg glomerulonephritis registry and the analysis performed. **B,** Schematic showing the cross section of a human kidney tissue with the localization of various cell types in different regions. Inset: cross section of a glomerulus(top) and tubulointerstitial region(bottom), and cell types therein. **C,** DAPI-stained images from kidney biopsies of control and ANCA-GN patient. The images show glomeruli and surrounding regions overlaid with cell boundaries determined using a cell segmentation algorithm. Localization of specific marker genes is indicated, with glomerular boundaries highlighted for clarity. **D,** UMAP projection of ∼3.2 million cells retained after quality control filtering. Cells are colored based on their annotated cell type. **E,** Stacked barplot showing the proportions of different cell types from the complete biopsy tissues across the four disease conditions. **F,** H&E images from control(left) and ANCA-GN (right) kidney biopsies showing a glomerulus and tubular interstitial region. For each region, the adjoining plots show the segmented cells color-coded according to their cell type annotation, illustrating cellular composition differences between conditions in different regions of the tissue. Cell type abbreviations: podocyte (podo), mesangial cell (MC), parietal epithelial cell (PEC), endothelial cell (EC), fibroblast (FIB), fibrotic mesangial cell (fib. MC), proximal tubule (PT), VSM/P: vascular smooth muscle cell/pericytes, distal convoluted tubule (DCT), connecting tubule (CNT), thick ascending limb of loop of Henle (TAL), ascending thin limb of loop of Henle (ATL), descending thin limb of loop of Henle (DTL), principal cell (PC) and intercalated cell (IC) of collecting duct, papillary tip epithelial cells abutting the calyx (PapE), neuronal cell (NEU), neutrophil (N), macrophages (MAC), monocytes (Mono), monocyte-derived cell (MDC), plasmacytoid dendritic cell (pDC), plasma cell (PL), conventional dendritic cell (cDC), cycling mononuclear phagocyte, NKC/T: natural killer cytotoxic T cell, cycNKC/T: cycling natural killer cytotoxic T cell.

**Table 1:** Baseline characteristics.

|  | **ANCA-GN (n=32)** | **SLE (n=19)** | **Anti-GBM (n=6)** |
| --- | --- | --- | --- |
| Age, median (IQR) | 55.5 (51 – 64.75) | 35 (29 – 42) | 57.5 (25.25 – 68.5) |
| Sex |  |  |  |
| Female, n (%) | 11 (34.37) | 14 (73.68) | 2 (33.33) |
| Male, n (%) | 21 (65.63) | 5 (26.32) | 4 (66.67) |
| Histology, n (%) | ANCA Renal Risk Score | Lupus-Nephritis Class | % crescentic or sclerotic glomeruli |
|  | low: 7 (24.14)<br>medium: 16 (55.17)<br>high: 6 (20.69) | Class III: 7 (38.89)<br>Class IV: 4 (22.22)<br>Class III+V: 5 (27.78)<br>Class IV+V: 2 (11.11) | <25: 0<br><50: 0<br><75: 2 (33.33)<br>≥75: 4 (66.67) |
| Laboratory values, median (IQR) |  |  |  |
| Creatinine (mg/dl) | 2.5 (1.78 – 3.84) | 0.9 (0.66 – 1.4) | 2.0 (1.93 – 6.81) |
| eGFR (ml/min) | 25.79 (13.77 – 36.4) | 89 (53 – 112) | 14 (8.92 – 31.28) |
| ACR (mg/g) | 1165 (378 - 2175) | 1100 (210 – 2130) | 1140 (325 - 11830) |
| Autoantibody levels (U/ml) | MPO: 88 (55.59 – 122.3)<br>PR3: 41 (1.47 – 145.3) | dsDNA: 55 (13 - 379) | Anti-GBM: 160 (29 – 765) |

### Panel design to identify immune cells, kidney cells and disease progression

Our first step was to design a comprehensive gene-panel for the identification of different kidney cells (Fig. 1B), immune cells and genes that may be important in the disease pathophysiology at the molecular level. We started with a detailed literature review to identify cell type-specific markers that would enable the comprehensive identification of all kidney and immune cells including subtype populations expected in our biopsy samples^5, 6, 7^. Next, we included genes that are (i) implicated in the pathogenesis of the cGN, (ii) essential for pathways involved in leukocyte infiltration and activation, (iii) associated with cytokine and chemokine signaling and inflammation for which we referred to the KEGG^8^, Reactome^9^ and Gene Ontology^10^ databases, and (iv) differentially expressed between normal and inflamed glomerular regions based on our recently published spatial transcriptomics data from patients with ANCA-GN and controls^11^. These steps resulted in the selection of 480 genes that were used for the high-resolution transcript analysis (Extended Data Table 1). Simultaneously, H&E staining was performed to visualize tissue morphology.

### Segmentation and classification of kidney and immune cells

To localize cells and determine their boundaries, we first segmented them using Baysor^12^, resulting in the identification of 3,218,210 filtered cells that passed our QC metric, with between 303,732 and 501,421 cells in each slide (Extended Data Fig. 1B-C). The cells expressed a median of 24 genes (control: 14, SLE: 25, ANCA-GN: 26 and anti-GBM: 27) and 36 median transcripts (control: 21, SLE: 41, ANCA-GN: 39, anti-GBM: 41) (Extended Data Fig. 1D). The median segmented cell diameter was approximately 7.7 μm^2^ (Extended Data Fig. 1E). We detected transcripts of cell-type-specific marker genes in both control and diseased glomeruli (Fig. 1C). These included markers for podocytes (*PODXL*), mesangial cells (*PDGFRB*), and PECs (*CLDN1*). In the tubulointerstitial regions, we found transcripts specific to proximal tubule cells (*GPX3*) and collecting duct cells (*AQP2*). Glomerulonephritis samples additionally showed expression of collagen (*COL1A1*) and immune cell markers, including those specific to monocytes and macrophages (*LYZ, CD163*) and T cells (*CD3D, CD3E, CD3G*) (Fig. 1C).

Subsequently, we used a logistic regression classifier to predict cell types with a median confidence >0.89 across cell types (Fig. 1D, Extended Data Fig. 2A). The cell types showed differential enrichment of cell type-specific marker genes (Extended Data Fig. 2B) and exhibited some of the expected differential abundance across diseases, such as the loss of podocytes and the increase of innate and adaptive immune cells in RPGN versus healthy controls (Fig. 1E). Exemplary localization of cell types within control and ANCA-GN glomerular and tubulointerstitial areas are presented in Fig. 1F together with the corresponding H&E-stained images, highlighting that glomerular and peri-glomerular regions are affected by cellular changes in disease. In summary, segmentation and classification resulted in a set of over 3 million spatially resolved renal and immune cells.

### Defining and analyzing the glomerular and peri-glomerular niches

Since in glomerulonephritis, the glomerulus and the surrounding tissue are most affected by inflammation and tissue destruction (Fig. 1F), we focused on these areas. Therefore, we needed a reliable annotation of the glomerular niches. To do this, we performed an automatic spatial domain annotation by employing NichePCA^13^ (Fig. 2A). The boundaries of the glomerular niches were subsequently extended by a 100 µm perimeter to define the peri-glomerular niches (Fig. 2B). We termed a region including a unique glomerulus and its associated peri-glomerular niche as a region of interest (ROI). Examples of identified glomerular and peri-glomerular niches are illustrated in Fig. 2C. Areas outside of glomerular and peri-glomerular niches were annotated as the tubulointerstitial domain. Fig. 2D illustrates selected examples of glomerular and periglomerular niches, and tubulointerstitial areas for each clinical condition with cells colored by their annotated types. Control and, to some extent, SLE samples showed healthy glomerular structures with a monolayer of PECs around the glomeruli and low numbers of immune cells. In the ANCA-GN glomerulus, an increased number of PECs formed a crescent structure inside the glomerulus. ANCA-GN and anti-GBM samples also showed increased accumulation of immune cells. A systematic comparison of the spatial accumulation of different cell types within the glomerular and periglomerular niches, and tubulointerstitial areas across various disease conditions is shown in Fig. 2E. The percentages of these cell types in individual patient samples, within the glomerular and periglomerular niches, are shown in Extended Data Fig. 3A and Extended Data Fig. 4A respectively. In the glomerulus, podocytes constituted a median of 30% of all cells in the control samples, which progressively decreased in SLE (25.9%), ANCA-GN (25%), and was the lowest in anti-GBM (23.8%) (Fig. 2E, Extended Data Fig. 3B). In contrast, the percentage of PECs in the glomerulus exhibited the opposite trend, showing an increase in the disease conditions. PECs constituted a median of 6% of all glomerular cells in control samples which increased to about 8% in SLE and more than 16% in ANCA-GN and anti-GBM samples (Fig. 2E, Extended Data Fig. 3B). Additionally, there was a decrease in the percentage of mesangial cells (control: 17.9%, SLE: 10.2%, ANCA-GN: 7.1%, anti-GBM: 4.3%) and a concomitant increase in fibrotic mesangial cells (control: 1.6%, SLE: 2%, ANCA-GN: 6.4%, anti-GBM: 8.8%) in the glomerulus. The fibrotic mesangial cells display a distinct fibrotic signature, characterized by increased production of ECM components such as collagens and fibronectin. A decline in the proportion of glomerular endothelial cells was also observed. Among immune cell types, a marked increase in innate immune cells was seen, particularly macrophages. They accounted for less than 0.5% of all glomerular cells in control samples which increased to median values of about 2% in SLE and ANCA-GN, and 7.8% in anti-GBM (Fig. 2E, Extended Data Fig. 3B). Other innate immune cells such as monocytes, MDCs, and neutrophils, were also increased within the glomerular niches. Notably, despite the increase in innate immune cells within the glomerular niche, there was negligible increase in adaptive immune cells such as T cells, natural killer cells (NKCs), B cells, or plasma cells in these regions (Fig. 2E, Extended Data Fig. 3B).

**Figure 2:**
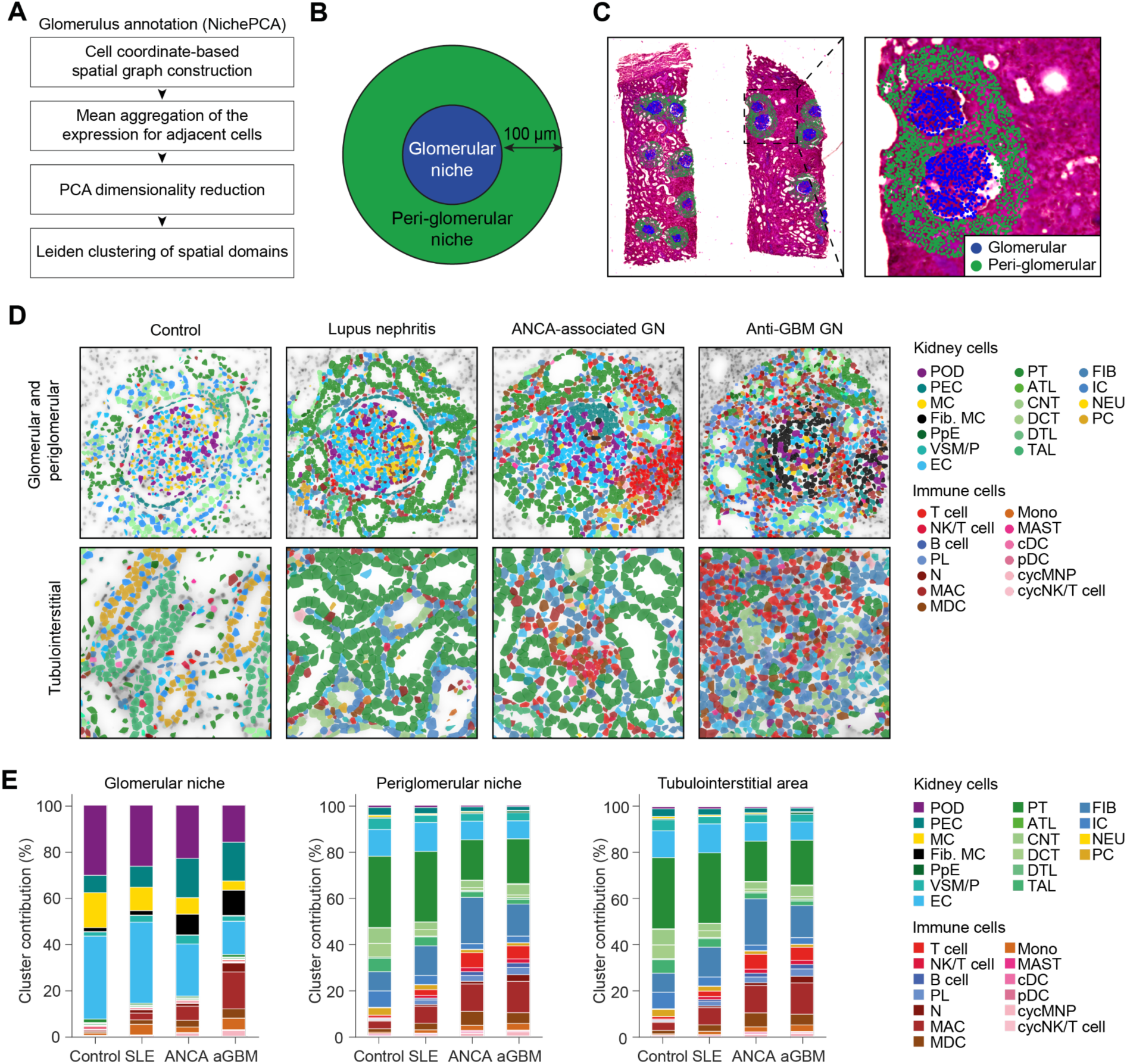
Identification and decoding the cellular composition of renal compartments by spatial transcript analysis in glomerulonephritis. **A,** Workflow of the NichePCA algorithm used for automated annotation and spatial mapping of glomeruli. **B,** Schematic illustration of a glomerulus and its periglomerular niche, defined by expanding the glomerular boundaries to include surrounding regions. **C,** Representative H&E image showing glomeruli (blue) as defined by NichePCA and their corresponding periglomerular regions (green). **D,** Representative images highlighting the spatial localization of various kidney and immune cell types. The top panels show the spatial localization of various kidney and immune cells within the glomerular and periglomerular regions, while the bottom panels depict the same in tubulointerstitial regions across the four disease conditions. **E.** Stacked bar plots showing the proportions of different cell types across the four disease conditions within three spatial niches: the glomerular niche (left), periglomerular niche (center), and tubulointerstitial areas (right).

In contrast, the periglomerular niches showed an increased accumulation of both innate and adaptive immune cells. For example, T cells, which in control samples constitute about 1.1% of all cells of the periglomerular domain, increased to a median of 2.5%, 4.8%, and 7.4% in SLE, ANCA-GN, and anti-GBM, respectively (Fig. 2E, Extended Data Fig. 4B). There was also a notable rise in the numbers of NKCs, B cells, and plasma cells in these regions and several innate immune cells in the periglomerular area (Fig. 2E, Extended Data Fig. 4B).

Taken together, we observed an increased accumulation of immune and fibrotic cell types in all three diseases, while the spatial accumulation of the immune cells showed distinct characteristics. The glomerular compartment in these diseases exhibited a selective accumulation of innate immune cells, with a notable absence of adaptive immune cell infiltration. However, the periglomerular domains exhibited an increased presence of both innate and adaptive immune cell types.

### Temporal trajectory of glomerular crescent formation

As outlined in the previous section, the different diseases showed qualitatively similar cellular changes across the three compartments, albeit with differing median quantities. These findings suggest that the path to glomerular crescent formation might be common across ANCA-GN, SLE, and anti-GBM, while temporal stages and the speed of progression might be distinct. In order to find common trajectories of disease progression across control and disease, we performed pseudo-time estimation of 769 combined unique ROIs (glomeruli plus periglomerular area). A single-path with progressive increase in pseudotime was observed across control and diseases, with ROIs from controls at the lowest pseudotime and from anti-GBM disease at the latest stage (Fig. 1A-B). To identify the molecular and cellular disease components of this trajectory, we compared the first principal component (PC1) of the ROIs to the pseudotime path, observing significant correlation (r=0.88). PC1 showed a cellular signature of crescent formation, including positive correlations with immune cells, PECs, and fibroblasts, while negatively correlating with podocytes and endothelial cells (Extended Data Fig. 5A-B). This suggests that pseudotime captures the common, progressive formation of crescents across the diseases, rather than the disease-specific progression time, which is partially captured in the other PCs.

To further understand the molecular and cellular characteristics of the spatio-temporal progression into glomerular crescents, we divided the pseudotime into four successive quadrants based on Jenks natural breaks optimization^14^ (0-0.36, 0.36-0.55, 0.55-0.77, and 0.77-1.0), each showing distinct distributions of conditions (Extended Data Fig. 5C). The first quadrant was dominated by control ROIs (80.34%). The second quadrant showed a mixed distribution, comprising control (42.77%), SLE (39.50%), ANCA-GN (16.57%) and anti-GBM (1.16%) ROIs. In contrast, the third and fourth quadrants were primarily characterized by ANCA-GN and anti-GBM conditions, with ANCA-GN predominating in quadrant 3 (70.31%) and anti-GBM being most prevalent in quadrant 4 (57.78%). We compared the gene expression profiles of quadrants to find upregulated marker genes for each quadrant in comparison to the other quadrants. Quadrants 1-2 showed upregulation of a limited number of genes, with a total of 31 and 35 differentially expressed genes (DEGs), respectively (Extended Data Fig. 5D). Quadrants 3 and 4, on the other hand, showed 177 and 137 DEGs. Among the top 10 genes enriched in the quadrants 3 and 4 were several immune-related cell type and fibrosis-associated genes (Immune: *CD79A, LYZ, S100A9*, and fibrosis: *FN1, LUM, TIMP1*) (Extended Data Fig. 5E)^15, 16^. Aggregating the top 5 most significantly enriched gene ontology (GO) biological processes per quadrant, we found several immune processes in quadrants 3 and 4 (Extended Data Fig. 5F). These two quadrants also showed a higher enrichment of several apoptotic and tissue damage processes, with a higher enrichment of these terms in quadrant 4 compared to quadrant 3 (Extended Data Fig 5G). Examples of ROIs within each quadrant are presented in Fig. 3C and indicate progressive crescent formation, qualitatively agreeing with the estimation of pseudotime trajectory.

**Figure 3:**
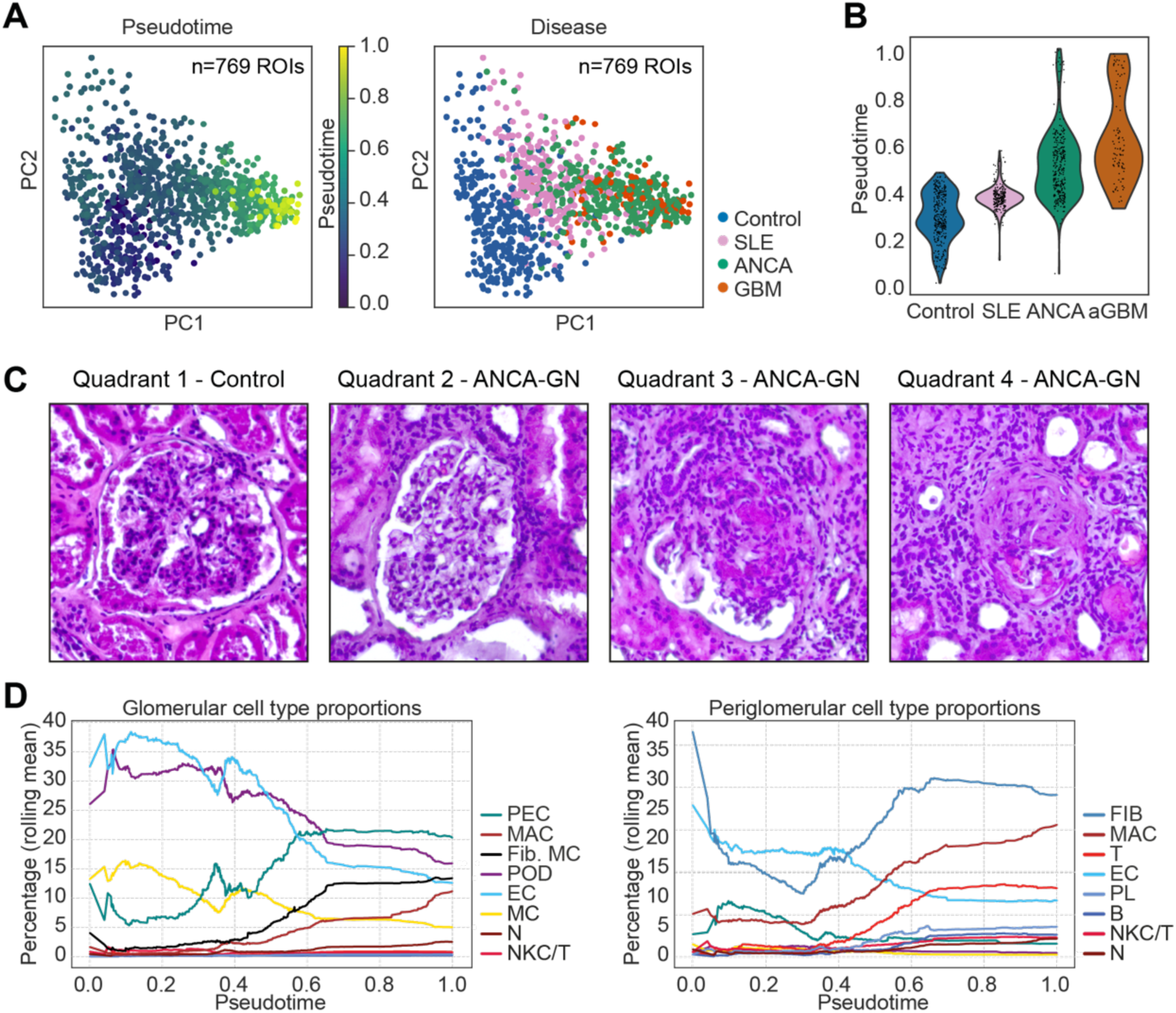
Trajectory of the immune cell ecosystem in glomerulonephritis. **A,** Principal Component Analysis (PCA) plot showing 769 unique regions of interest (ROIs), each comprising a distinct glomerulus and its associated periglomerular area. ROIs are colored by pseudotime (left) and by disease condition (right). **B,** Distribution of pseudotime values assigned to the ROIs across the four disease conditions. **C,** Representative H&E images showing glomeruli from each of the four pseudotime quadrants, highlighting morphological variations. **D,** Change in the percentage distribution of different cell types within glomerular regions (left) and periglomerular regions (right) across pseudotime.

These results indicate that SLE, ANCA-GN, and anti-GBM share a common cellular and molecular trajectory of crescent formation, which might contain shared inter- and intra-cellular signaling pathways. While the data also indicated disease-specific temporal changes, the common crescentic path dominated both the pseudotime and PC-based analyses.

### Inter-cellular signaling of glomerular crescent formation

To elucidate the intercellular signaling pathways that underlie the pan-RPGN temporal development of glomerulonephritis, we performed interactome analyses using CellChat v2, which provides a framework for inferring interaction probabilities between cell-types by integrating the ligand-receptor expression data, as well as the spatial localization of these cell populations^17^. We focused on the regions of biggest change, the glomerular and periglomerular regions. In general, cGN samples were found to have a higher number of cell-cell interactions as compared to those in control samples. A comparison of the cell-interaction networks between cGN and control samples revealed that PECs exhibited increased interactions with neighboring cell types in glomerulonephritis (Fig. 4A). Given the increased signaling activity originating from PECs and the central role of PECs in crescent formation, we next investigated which pathways result in activation of PECs. To this end, we filtered all significant cell-cell interactions in cGN, to extract the signaling interactions that are directed towards PECs as the target cell. This analysis revealed that PDGF, TGF-β, FASL and SLIT pathways showed significant activity towards PECs (Fig. 4B). The primary source of PDGF ligands to PECs were endothelial cells followed by mesangial cells, vascular smooth muscle cells/pericytes, PECs and macrophages. TGF-β was the other key signaling pathway targeting PECs, with the most prominent source of TGF-β ligands being the fibrotic mesangial cells. Additionally, PECs received TGF-β ligands from endothelial cells, podocytes, PECs, fibroblasts, and different innate immune cell types (macrophages, monocytes) as well as adaptive immune cells (T cells, NKC/T and B cells) (Fig. 4B).

**Figure 4:**
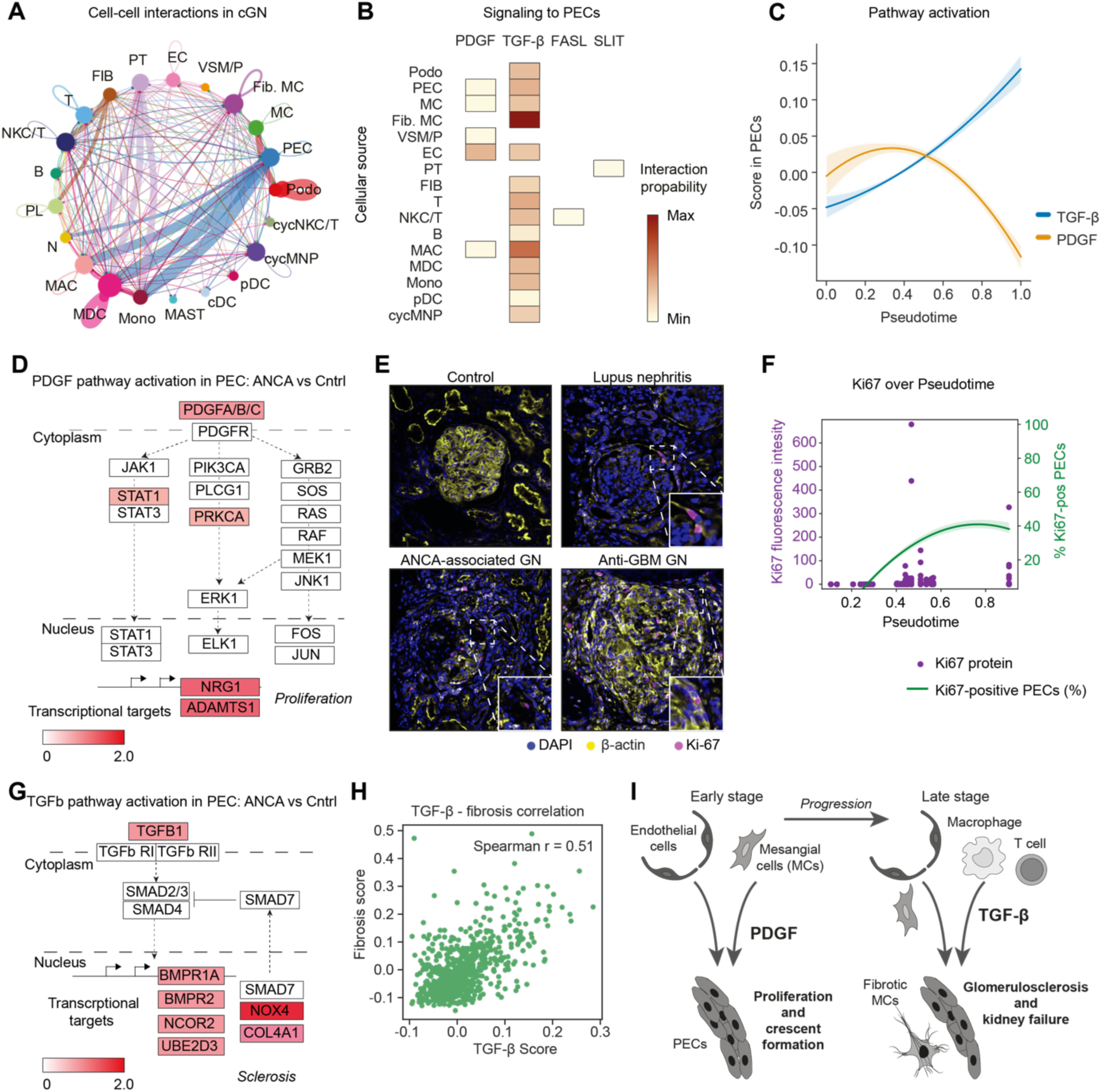
Mapping the immune-epithelial cell interactions in glomerular crescent formation. **A,** Circleplot showing cell-cell interactions in crescentic glomerulonephritis (cGN) compared to control samples. The interaction edges are colored the same as the source cell-type while edge weights are proportional to the interaction strength. **B,** Heatmap showing the signaling pathways that have increased activity directed towards PECs in cGN, and the source cell types producing the corresponding ligands, colored by the interaction probability. **C,** Scores for PDGF and TGF-β pathway activation in PECs plotted along pseudotime. **D,** Schematic representation of the PDGF signaling pathway, highlighting the genes upregulated in PECs in a snRNA-seq dataset from ANCA-GN patient samples (n=6) over controls (n=3). **E,** Multiplexed protein staining on a Xenium slide showing overexpression of Ki67 in cGN samples. **F.** Expression of Ki67 protein in PECs along pseudotime. The primary axis corresponds to fluroscence intensity values of Ki67. The secondary axis curve shows change in percentage of Ki67-postive cells per glomerulus. **G,** Schematic representation of the TGF-β signaling pathway, highlighting the genes from this pathway upregulated in PECs in a snRNA-seq dataset from ANCA-GN patient samples (n=6) over controls (n=3). **H,** Correlation of TGF-β pathway score with Fibrosis score in PECs. **I,** Schematic illustrating the role of PDGF and TGF-β mediated activation of PECs in crescent formation and glomerulosclerosis in cGN.

### PDGF induces PEC proliferation and crescent formation

We next investigated the dynamics and effects of early PDGF activation in PECs in glomerulonephritis. Therefore, we quantified the activation levels of the PDGF pathway in PECs by creating a pathway-specific score using gene sets that capture the level of activation of the pathway. We used the Reactome gene set for PDGFR signaling in disease, available within the curated gene sets collection (C2) from the Molecular Signatures Database (MSigDB)^18^. We plotted this PDGFR score in PECs against pseudotime derived in the previous analysis (Fig. 4 C). The PDGF score exhibited an increase at early to medium pseudotime stages, peaking approximately at 0.35 (Q2) along the pseudotime.

To validate the activation of PDGF pathways observed in PECs within the spatial transcriptomics dataset using an independent orthogonal dataset with higher transcript resolution per cell, we generated a single nuclear transcriptome sequencing (snRNA-seq) dataset from kidney biopsies of ANCA-GN patients (n=6) and controls (healthy part of tumor nephrectomies, n=3), resulting in a total of 112,560 nuclei after QC filtering. The nuclei expressed a median of 899 genes and 2128 median transcripts (Extended Data Fig. 7A). After preprocessing and clustering, cell types were predicted using a logistic regression classifier, achieving a median confidence >0.98 across the cell types (Extended Data Fig. 7B). The predicted cell types displayed differential enrichment of marker genes specific to each cell type (Extended Data Fig. 7B). Subsequently, we identified genes that were differentially expressed between PECs from ANCA-GN and control samples (Wilcoxon test, adjusted p-value < 0.05). Genes associated with the PDGF pathway were found to be upregulated in PECs from ANCA-GN compared to control samples, further confirming the activation of these pathways in ANCA-GN-related pathology (Fig. 4D). Genes transcriptionally upregulated downstream of the PDGF signaling pathway included those involved in regulating proliferation, such as *NRG1*^19^, and genes associated with fibrosis such as *ADAMTS1*^20^ (Fig. 4D).

To investigate if high PDGF signaling activity during the early stages of crescent formation indeed results in proliferation of PECs, we investigated the presence of Ki-67 protein along the pseudo-time. To this end, we combined Xenium transcript analysis with Akoya Phenocycler-based spatial proteomics of the same slide for 8 samples (2 controls, 3 ANCA-GN, 2 SLE and 1 anti-GBM disease, Fig. 4E). To align spatial transcriptomic and proteomic images, we registered the Akoya Phenocycler and Xenium DAPI signals using VALIS ^21^. This registration was subsequently applied to all other protein stains. Following registration, Ki-67 intensity levels were assigned to PECs within all ROIs based on the cell boundaries obtained by our cell segmentation (Extended Data Fig. 8). Leveraging the two datasets generated by the independent analyses of RNA and protein measurements from the same histological slide, we were able to identify PECs based on their transcriptional profile, align glomeruli according to the pseudo-time, and determine cell proliferation by Ki-67 quantification. These analyses confirmed that PEC proliferation assessed via Ki-67 fluorescence and percentage of Ki-67 positive PECs per glomerulus increased early along the pseudotime after PDGF activation and followed similar dynamics (Fig. 4F) as PDGF signaling in PECs (Fig. 4C).

### TGF-β signaling results in PEC- and mesangial cell-mediated glomerular sclerosis

After clarifying the role of early PDGF signaling on PECs, we next wanted to determine the functional effects of TGF-β signaling on PECs in the phase transition from inflammatory cellular crescents to irreversible fibrotic crescents. We used the human gene set of the TGF-β signaling signature under the Hallmark gene sets collection (H) from MSigDB^18^. When changes in pathway activation were plotted along the crescent development, the TGF-β score showed a continuous increase over the pseudo-time (Fig. 4C). For TGF-β, snRNA-seq DEGs from ANCA versus control PECs revealed upregulation of TGF-β pathway genes that play crucial roles in fibrosis, including COL4A1 and NOX4 (Fig. 4G). COL4A1 encodes type IV collagen, a key component of the ECM. In fibrotic conditions, increased deposition of type IV collagen contributes to ECM stiffening and tissue remodeling^22^. NOX4 is a major source of reactive oxygen species (ROS) and promotes oxidative stress which drives fibrotic processes by activating myofibroblasts and enhancing ECM deposition^23, 24^. The role of NOX4 in renal fibrosis is complex and it has also been shown to have a protective role^25^. Other genes activated by the TGF-β pathway are BMPR1A and BMPR2, which are receptors for BMP4, a ligand within the TGF-β superfamily, and NCOR2, a transcriptional corepressor that modulates fibrosis by regulating TGF-β target genes^26^.

Finally, we aimed at understanding if TGF-β signaling in PECs indeed could result in glomerulosclerosis in human glomerulonephritis. Therefore, we investigated the TGF-β signaling score and a score that includes markers for fibrosis (Fig. 4H) and found a significant correlation. These findings support the hypothesis, that first PDGF from endothelial cells and mesangial cells induces cell proliferation of PECs and glomerular crescent formation, and later in the disease immune and kidney cell derived TGF-β results in PEC- and fibrotic mesangial cell-mediated glomerular sclerosis (Figure 4I).

## Discussion

Rapidly progressive glomerulonephritis (RPGN) is characterized by immune cell infiltration into the kidney and glomerular crescent formation. The lack of knowledge about the immune and kidney cells as well as their interactions, leading to glomerular damage and disease progression, has hindered the development of target therapies. Here, we report a spatial high-resolution transcriptomic atlas of kidney biopsies from RPGN patients, deciphering the cell composition and their communications involved in sequence of glomerular crescent formation. RPGN is a group of severe, rapidly progressing autoimmune diseases marked by the formation of cellular crescents in Bowman’s space, leading to severe glomerular injury and loss of kidney function. The most common causes of RPGN include ANCA-GN, SLE and anti-GBM disease, in which the pathology is triggered by an intense inflammatory reaction within the glomerulus and often the periglomerular region^27^. Experimental GN models demonstrated that the recruitment of immune cells and the production of various inflammatory mediators^28^ lead to the rupture of glomerular capillaries and the activation and proliferation of parietal epithelial cells (PECs), which together with infiltrated leukocytes form the characteristic glomerular crescent^29, 30, 31, 32^.

Recent single-cell and spatial transcriptome analyses in renal biopsy samples improved our understanding of kidney immunity in RPGN patients by identifying inflamed regions, leukocytes and predominant cytokine signaling in the tissue^11, 33, 34, 35^. However, identification of immune cell subtypes and their precise localization in the context of crescentic GN was so far not possible due to the low resolution of the spatial approach^11^. By using high-resolution spatial transcriptomics technologies^2, 3^, we precisely define immune and kidney cells within the glomerular, periglomerular, and interstitial regions in a large cohort of RPGN patients. We observed a notable enrichment of innate immune cells such as monocyte-derived cells and macrophages within the glomeruli in all three cGN subtypes. In contrast, adaptive immune cells were predominantly found in the periglomerular regions.

Additionally, our investigation of cell-cell interactions among all cell types in the glomerular and periglomerular compartments, encompassing both structural kidney cell types and immune cells highlighted increased activation of PECs, driven by PDGF signaling primarily from endothelial cells, mesangial cells and vascular smooth muscle cells/pericytes. Pseudo-time analysis of our spatial transcriptomic data combined with whole kidney snRNA-seq data formation in the “inflammatory” phase of crescentic GN. The platelet-derived growth factor (PDGF) family comprises four ligand isoforms (PDGF-A, -B, -C, and -D) and two receptor chains, PDGFR-α and PDGFR-β, which are either constitutively or inducibly expressed across most renal cell types^36^, including PECs^37^. In the context of kidney diseases, *de novo* expression or upregulation of PDGF isoforms and their receptors has been reported in various rodent models of renal injury, as well as in corresponding human renal pathologies. Binding of ligands to PDGF receptors induces auto-phosphorylation within the cytoplasmic tyrosine kinase domain of the receptor, thereby facilitating the recruitment of adapter proteins with SH2 and SH3 domains, further activating downstream signaling pathways, including the JAK/STAT, PI3K/AKT, PLC-γ, and RAS/MAPK pathways, which induce gene expression supporting proliferation and migration of the receptor expressing cells as well as fibrosis^36^. While the signaling cascade is not regulated at the transcriptional level, we see downstream effects such as upregulation NRG1 and proliferation in PECs.

The factors which promote the transition from the inflammatory phase of crescentic GN, defined by cellular crescents (which can be reversible), to an irreversible fibrotic phase with fibrous crescents and glomerulosclerosis is largely unknown. Our finding that TGF-β downstream signaling in PEC is predominantly detectable at later stages of the pseudo-time analysis is therefore of interest. TGF-β is known as a central driver of tissue fibrosis, promoting extracellular matrix (ECM) protein synthesis, inhibiting matrix degradation, and inducing cell transformations that lead to glomerulosclerosis and tubulointerstitial fibrosis. Elevated TGF-β levels are strongly associated with progression in cGN^38, 39, 40^. TGF-β induces fibrosis through both SMAD-dependent and independent pathways. TGF-β activation of fibroblasts and epithelial-to-mesenchymal transition (EMT) further amplifies fibrotic processes^41, 42, 43^. Various approaches to inhibit TGF-β have been explored, including neutralizing antibodies, receptor inhibitors, siRNA, and small molecule inhibitors. Clinical trials with anti-TGF-β agents such as LY2382770 and fresolimumab in fibrotic kidney diseases have yielded mixed results^44, 45, 46^. The failure of drug response is attributed to the pleiotropic nature of TGF-β in both immune regulation and kidney fibrosis. However, our data suggest that TGF-β plays a more dominant role in the later stages of disease progression, so interventions at this stage may be more promising for patients with advanced disease.

These analyses define the cell infiltration and cell communications in the progression of and the identification of important drivers of glomerulosclerosis paves the way for future therapies interfering with crescent formation and blocking phase transition to fibrosis. The spatial high-resolution atlas of human glomerulonephritis provided here will also serve as a key resource for studies that investigate cell infiltration, communication and pathways involved in these diseases.

## Methods

### Study design and patient inclusion

This study was designed to analyze immune cells and parenchymal kidney cells as well as transcriptional profiles of inflamed glomeruli in the kidney tissue of patients with rapidly progressive glomerulonephritis and in homeostatic kidney tissue. All patients with glomerulonephritis were included in the Hamburg Glomerulonephritis Registry. Health kidney tissue was obtained from unaffected kidney parts of tumor nephrectomies. All subjects provided informed consent in accordance to the CARE guidelines and the ethical principles stated in the Declaration of Helsinki. Information about the sex of the patients is provided in Table 1. Biological sex and self-reported sex were identical. Sex- and gender-based analyses were not performed in this study. This study was approved by the Institutional Reviewing Board (IRB) of the University Medical Center Hamburg-Eppendorf and the *Ethik-Kommission der Ärztekammer Hamburg* (local ethics committee of the chamber of physicians in Hamburg; licenses PV4806, PV5026, and PV5822).

### High-dimensional spatial data generation

For performing spatial analysis, human kidney samples from FFPE embedded tissue were used. 5 µm thick tissue sections were placed on Xenium slides and processed according to manufacturer’s instructions (10x Genomics, Pleasanton, CA, USA). Subsequently, HE staining was performed and whole slide images were obtained on a Leica DMi8 system using the Leica Application Suite X software Version 3.7.4.23463 software (Leica, Wetzlar, Germany)^11^. In case of protein analysis, protein staining and image acquisition was performed on a PhenoCycler-Fusion system (Akoya Biosciences, Marlborough, MA, USA) according to manufacturer instructions before HE staining.

### Data processing

The default nuclei-expansion based segmentation from Xenium was found to be suboptimal with respect to the cell area and purity of cell types in terms of marker genes, an observation similar to Salas et al., 2023^47^. To find the optimal cell segmentation for the Xenium data, the default segmentation was refined using Baysor (version 0.6.2)^12^ with confidence in prior probability set to 0.5. To exclude artifacts, cells from the tissue regions that were excessively blurred or showed potential noisy regions (such as folded and overlapping tissues) were removed. Additionally, the cells belonging to spots outside of the biopsy were excluded. Finally, cells that expressed less than five unique genes were removed.

Published single-cell reference data from the kidney^6^ was used to train a machine learning classifier. 472 genes in the reference data were common with the gene panel. The reference data and processed Xenium data was subsetted to include only the genes that were common between the both. The resulting datasets were log-normalized using *scanpy.pp.normalize_total* and *scanpy.pp.log1p* methods from Scanpy (version 1.10.1). The target_sum was set to 1,000. LogisticRegressionCV class from scikit-learn (version 1.3.1) was used to train for the classification task with input being the normalized count matrix of the reference data and target being the cell type labels. The trained model was applied to the similarly normalized Xenium data and the labels were transferred to the original processed object.

### Defining the glomerular domain

To identify the glomerular niches based on the spatial transcriptome data, we used NichePCA^13^. Briefly, NichePCA constructs a spatial graph based on the spatial coordinates of cells and then calculates the mean aggregation of gene expressions for adjacent cells. Next, principal component analysis (PCA) is performed to reduce the dimensionality of the aggregated expressions, and finally Leiden clustering is used to identify the spatial domains and here resulted in the definition of glomeruli (Fig. 2A). Further, cells showing fibrotic signature inside the glomeruli regions were labelled as fibrotic mesangial cells.

### Cell-cell interaction analysis

We used CellChat (version 2.1.2)^17^ to analyze cell-cell interactions which utilizes the x and y coordinates of the cells available from spatial transcriptomic datasets to infer cell-cell communication taking into account the spatial proximity of cells to infer probability of communication between them. Since the 63 biopsy samples in this analysis were distributed across 8 Slides, we added offset values to the *x*-coordinates of cells to all slides except the first, thereby generating unique *x* and *y* coordinates for each slide to ensure that spatial interactions were only inferred among cells on the same slide.

Next, we generated separate CellChat objects for each of the four categories of samples (Cntrl, SLE, ANCA, anti-GBM) using the *createCellChat* function. We then set the ligand-receptor interaction database to include only “Secreted Signaling” using the *subsetDB* function. Next, we preprocessed the expression data for cell-cell communication analysis using the *identifyOverExpressedGenes* and *identifyOverExpressedInteractions* functions. To compute the communication probability and infer cellular communication network, we used the *computeCommunProb* function with the *truncatedMean* method, applying a trim value of 0.1 to compute the average gene expression, and set the parameters *interaction.range* to 250 and *contact.range* to 10. The cell-cell communication at a signaling pathway level was inferred using the *computeCommunProbPathway* function and aggregated cell-cell communication network were combined using the *aggregateNet* function. Next, we calculated the difference of the interaction weight matrices for the cGN conditions over control to visualize the difference seen in the cGN conditions with respect to the communication network that exists in control (Fig. 4A). The change in interaction weights for each of the three diseases individually over control is shown in Extended Data Fig. 6A.

In order to investigate all incoming signaling directed toward PECs, we used the subsetCommunication function to obtain a data frame of significant cell-cell interactions from each CellChat object. This data frame contains detailed information on significant inferred cell-cell communications. We then combined the data from the control and disease conditions into a comprehensive data frame representing all interactions. This data frame was then used for visualizing all incoming signaling directed toward PECs, which primarily included PDGF and TGF-β pathway interactions (Fig. 4B). The signals for each of the disease types individually in shown in Extended Data Fig. 6 B.

### Data integration / Integrating transcript and protein data / HE staining

The transcriptome data was used as the fixed reference, while the proteomics and HE staining data were transformed to align with it. In particular, for the transcriptomics and proteomics data, the DAPI staining was used, and the other protein channels were transformed accordingly. For image registration, we cropped the *.tiff* image files around each sample and applied the non-rigid registration workflow available in VALIS v1.1.0. ^21^. The quality of registration was assessed by visualizing stacked images in FIJI v 1.54. (https://doi.org/10.5281/zenodo.5256255). Following registration, protein expression levels were assigned to individual cells. This was done by summing the pixel intensities of proteins located within the cell boundaries (using shapely python package), as defined by the cell segmentation algorithm.

### Pseudotime / trajectory analysis

The raw counts of each ROI (unique glomerular + peri-glomerular areas) were summed to create pseudo bulk ROIs. The resulting counts were normalized to sum to 10,000 and log normalized using preprocessing.normalize_total and preprocessing.log1p methods from Scanpy (version 1.10.1)^48^. Pseudotime computation was performed using diffusion pseudotime method^49^ with a starting point to be a random control ROI. Pseudotime was further divided into four clusters termed “quadrants” using Jenkins natural breaks optimization using python package jenkspy (v0.4.1)^14^.

### Single nucRNA-sequencing

For single nucRNA-sequencing, snap frozen biopsies were thawed and chopped using a fresh razor blade. After adding 200 µl lysis buffer (1 mM DTT, 0.1% Triton X-100, 0.2 U/µl SUPERaseIN RNAse Inhibitor and 0.4 U/µl RNasin® Plus Ribonuclease Inhibitor in Nuclei PURE Lysis Buffer), the tissue homogenized 60 times using a pestle on ice. Next, the lysate was supplemented with additional 3.8 ml lysis buffer and incubated on ice for 25 min. Lysis efficiency was assessed using an optical microscope. After incubation, a 30 µm strainer was used to filter the lysate and washed with 1 ml lysis buffer. The lysate was centrifuged at 500 g (5 min, 4°C) and the supernatant was discarded. 5 ml washing buffer (0.01% BSA, 0.4 U/µl SUPERaseIN RNAse Inhibitor and 0.4 U/µl RNasin® Plus Ribonuclease Inhibitor in PBS) were added to the pellet and transferred to a falcon tube coated with 0.1% BSA in PBS and centrifuged (500 g, 5 min, 4°C). Resuspension buffer (1% BSA and 0.4 U/µl RNasin® Plus Ribonuclease Inhibitor in PBS) was added to the pellet. 40,000 nuclei were loaded onto a Chip G for GEM generation using the Chromium Single Cell 3’ v3.1 kit (10X Genomics). Reverse transcription, barcoding, complementary DNA amplification and purification for library preparation were performed as specified in the manufacturer’s protocol. Sequencing was performed on a NovaSeq 6000 platform (Illumina) at a target read depth of 25,000 by Novogene.

Cell Ranger software (v7.1.0, 10x Genomics) was used to process cellular barcodes and align reads to the GRCh38-2020-A reference genome. Quality control and preprocessing was performed using Seurat (v4.9.9) and R (v4.3.1). Nuclei expressing fewer than 200 genes or more than 2500 genes were excluded. Additionally, low-quality cells with over 5% mitochondrial genes were filtered out. The raw count data were normalized to 10,000 reads per nucleus, log-transformed (log1p), batch corrected and integrated with Seurat using the 2000 most highly variable genes. Clustering was performed using the Louvain algorithm with resolution 0.8 and cell types were annotated using a logistic regression classifier.

## Acknowledgements

We thank the Single Cell Sequencing and the Cytometry and Cell Sorting core facilities at the University Medical Center Hamburg-Eppendorf for their support. This study was supported by grants from the *Deutsche Forschungsgemeinschaft*, DFG (SFB 1192 to V.P., T.B.H., U.P, S.B., C.F.K.; KR 3483/3-1 to C.F.K.). RK was supported by DFG FOR 5068. B.Y. was supported by the Federal Ministry of Education and Research (BMBF) as part of the German Center for Child and Adolescent Health (DZKJ).

## Author contributions

U.P., S.B. and C.F.K. conceptualized the study. Z.S., R.K., and B.Y. conducted data analysis. N.S., S.J.-S., V.S., A.P., A.K., A.B. and H.-J.P. processed samples. N.S., S.J.-S., V.S., A.H., T.G.-S. acquired data. D.P.S. processed data. Z.S., R.K., B.Y., U.P. and C.F.K. designed figures. Z.S., R.K., B.Y., S.B., U.P. and C.F.K. wrote the manuscript. J.E., M.H., U.O.W., E.H., T.W., T.B.H., U.P. and C.F.K. provided patient samples. V.P., U.P., S.B., C.F.K. supervised the study. All authors approved the final manuscript.

## Materials & Correspondence

Correspondence and material requests should be addressed to U.P., S.B. or C.F.K.

## Extended data figure legends

**Extended Data Fig. 1: Quality control metrics and cellular characteristics of Xenium spatial transcriptomics data. A,** Number of samples belonging to control and disease conditions across the 8 Xenium slides. **B-C,** Number (B) and distribution (C) of cells in each slide and condition after filtering for quality control. **D,** Boxplot showing distribution of number of unique transcripts (left) and number of unique genes (right) detected in each cell. **E,** Boxplot showing distribution of segmented cell-areas per cell type. The median cell area and approximate diameter overall cells are indicated by a dashed line. Boxplots in panels D and E show the median (middle horizontal line), interquartile range (box), Tukey-style whiskers (lines beyond the box), outliers (data points beyond 1.5*interquartile or below -1.5*interquartile).

**Extended Data Fig. 2: Cell type classification metrics and marker gene expression profiles. A,** Matrix showing the classifier confidence (median, mean and standard deviation in prediction of the cell types. **B,** Heatmap showing the expression of cell type marker genes in annotated cell types: podocyte (POD), mesangial cell (MC), parietal epithelial cell (PEC), endothelial cell (EC), fibroblast (FIB), fibrotic mesangial cell (fibrotic MC), proximal tubule (PT), VSM/P: vascular smooth muscle cell/pericytes, distal convoluted tubule (DCT), connecting tubule (CNT), thick ascending limb of loop of Henle (TAL), ascending thin limb of loop of Henle (ATL), descending thin limb of loop of Henle (DTL), principal cell (PC) and intercalated cell (IC) of collecting duct, papillary tip epithelial cells abutting the calyx (PapE), neuronal cell (NEU), neutrophil (N), macrophages (MAC), monocytes (Mono), monocyte-derived cell (MDC), plasmacytoid dendritic cell (pDC), plasma cell (PL), conventional dendritic cell (cDC), cycling mononuclear phagocyte, NKC/T: natural killer cytotoxic T cell, cycNKC/T: cycling natural killer cytotoxic T cell.

**Extended Data Fig. 3: Cell type composition analysis of glomerular regions. A,** Stacked bar plot showing the cell type proportions for individual glomerular regions across control and disease conditions. **B,** Boxplots showing comparisons of the cell type proportions in glomerular regions between control, SLE, ANCA-GN, and anti-GBM conditions.

**Extended Data Fig. 4: Cell type composition analysis of periglomerular regions. A,** Stacked bar plot showing the cell type proportions for individual periglomerular regions across control and disease conditions. **B,** Boxplots comparing cell type proportions in periglomerular regions between control, SLE, ANCA-GN, and anti-GBM conditions.

**Extended Data Fig. 5: Relationship between principal component analysis and pseudotime. A,** Scatter plots showing linear associations between the first four principal components (PC1-4) and pseudotime for ROIs, with Pearson correlation coefficients (r) and p-values. **B,** Heatmap showing Pearson correlations between cell type abundances and principal components (PC1-4) in ROIs. **C,** Matrix showing the percentage distribution of conditions across the pseudotime quadrant. **D,** Bar plot showing the number of markers identified in each pseudotime quadrant. **E,** Heatmap displaying the mean expression of top 10 markers for each pseudotime quadrant, with immune and crescent scores shown below. **F,** Heatmaps showing top enriched GO biological process terms for each pseudotime quadrant. 5 most significant terms were selected for each quadrant. **G,** Heatmap showing enriched terms specifically related to fibrosis, extracellular matrix (ECM) remodeling, and tissue damage across pseudotime quadrants.

**Extended Data Fig. 6: Cell-cell communication networks and PEC-specific signaling in glomerular diseases. A,** Circular network diagrams depicting intercellular communication patterns in SLE, ANCA-GN, and anti-GBM. Nodes represent different cell types and edges indicate communication strength between cell pairs, with edge thickness proportional to interaction probability. **B,** Heatmap showing disease-specific cell-type sources and their signaling pathways targeting PECs across SLE, ANCA-GN, and anti-GBM conditions.

**Extended Data Fig. 7: Single nuclear mRNA sequencing (snRNA-seq) QC and cell type annotation. A,** Violin plots show quality control metrics, distribution of number of genes, total counts, percentage of mitochondrial and ribosomal genes across samples. Barplots show the total number of nuclei per sample. **B,** Matrix showing the classifier confidence (median, mean and standard deviation in prediction of the cell types. **C,** Heatmap showing the expression of cell type marker genes in annotated cell types.

**Extended Data Fig. 8: Aligning spatial transcriptomic and proteomic data. A,** The DAPI stainings of spatial transcriptomic and proteomic data were registered using the VALIS python package. **B,** This registration was applied to all other protein stains. **C,** DAPI staining of the spatial transcriptomic data for a selected SLE glomeruli. The red polygons represent boundaries identified for PECs. **D,** DAPI staining of the spatial proteomic data after registration to the DAPI staining of the spatial transcriptomic data. **E,** Ki-67 protein image after registration. **F,** PECs color-coded based on the cumulative pixel intensities of Ki-67 within their boundaries. **G,** CD44 protein image after registration. **H,** PECs color-coded based on the cumulative pixel intensities of CD44 within their boundaries.

## Extended data table legend

**Extended Data Table 1: Gene-panel designed for the identification of different kidney cells, immune cells and glomerulonephritis driving genes.**

