## Extended Data for "Spatio-temporal interaction of immune and renal cells determines glomerular crescent formation in autoimmune kidney disease"

Extended Data Fig 1

A

Number of biopsies in each Xenium slide

| | Control | SLE | ANCA | aGBM |
|---|---|---|---|---|
| 0011216 | | 3 | 4 | 1 |
| 0011284 | 1 | 2 | 4 | 1 |
| 0011287 | | 3 | 5 | |
| 0011546 | 2 | 2 | 3 | 1 |
| 0011695 | | 2 | 5 | 1 |
| 0011707 | | 3 | 4 | 1 |
| 0011762 | 2 | 2 | 3 | 1 |
| 0018775 | 1 | 2 | 4 | |

B

Number of cells in each slide and condition

| | Control | SLE | ANCA | aGBM | Total |
|---|---|---|---|---|---|
| 0011216 | | 151,011 | 129,585 | 220,825 | 501,421 |
| 0011284 | 52,394 | 63,985 | 195,419 | 45,694 | 357,492 |
| 0011287 | | 128,889 | 315,988 | | 444,877 |
| 0011546 | 119,946 | 82,456 | 140,280 | 65,073 | 407,755 |
| 0011695 | | 134,869 | 251,374 | 40,523 | 426,766 |
| 0011707 | | 157,540 | 180,432 | 58,421 | 396,393 |
| 0011762 | 121,729 | 14,574 | 135,230 | 32,199 | 303,732 |
| 0018775 | 90,146 | 64,904 | 224,724 | | 379,774 |
| Total | 384,215 | 798,228 | 1,573,032 | 462,735 | 3,218,210 |

C

Proportions of cells in each slide and condition

| | Control | SLE | ANCA | aGBM |
|---|---|---|---|---|
| 0011216 | | 0.19 | 0.08 | 0.48 |
| 0011284 | 0.14 | 0.08 | 0.12 | 0.10 |
| 0011287 | | 0.16 | 0.20 | |
| 0011546 | 0.31 | 0.10 | 0.09 | 0.14 |
| 0011695 | | 0.17 | 0.16 | 0.09 |
| 0011707 | | 0.20 | 0.11 | 0.13 |
| 0011762 | 0.32 | 0.02 | 0.09 | 0.07 |
| 0018775 | 0.23 | 0.08 | 0.14 | |

D

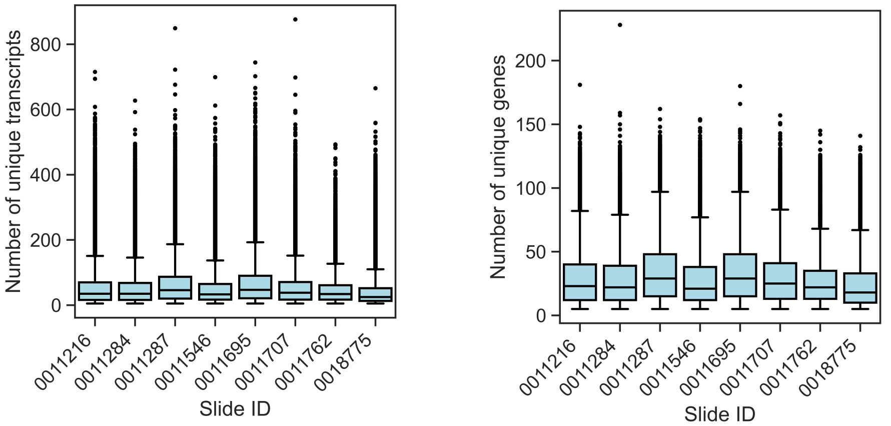


E

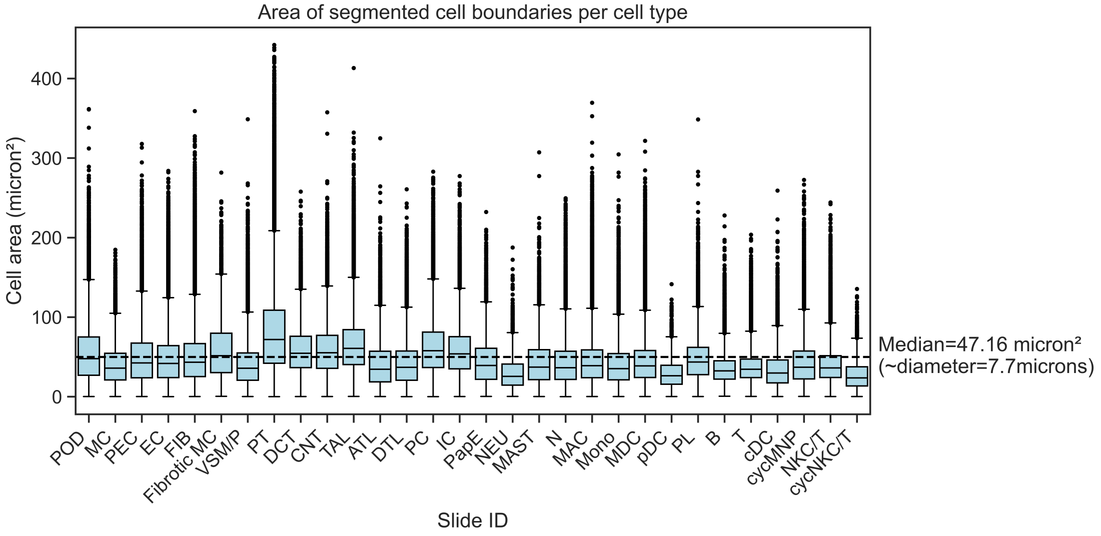

Extended Data Fig 2

A

Cell type confidence

| | Median | Mean | Standard deviation |
|---|---|---|---|
| POD | 1.00 | 0.95 | 0.12 |
| PEC | 1.00 | 0.93 | 0.14 |
| MC | 1.00 | 0.92 | 0.15 |
| Fibrotic MC | 1.00 | 0.94 | 0.12 |
| PapE | 0.95 | 0.86 | 0.17 |
| VSM/P | 1.00 | 0.93 | 0.14 |
| EC | 1.00 | 0.94 | 0.12 |
| PT | 1.00 | 0.97 | 0.09 |
| ATL | 0.97 | 0.88 | 0.17 |
| CNT | 1.00 | 0.93 | 0.13 |
| DCT | 1.00 | 0.96 | 0.10 |
| DTL | 0.99 | 0.90 | 0.16 |
| TAL | 1.00 | 0.96 | 0.11 |
| FIB | 1.00 | 0.98 | 0.08 |
| IC | 1.00 | 0.97 | 0.09 |
| NEU | 1.00 | 0.92 | 0.15 |
| PC | 1.00 | 0.95 | 0.12 |
| T | 1.00 | 0.95 | 0.12 |
| NKC/T | 1.00 | 0.91 | 0.15 |
| B | 1.00 | 0.94 | 0.14 |
| PL | 1.00 | 0.94 | 0.13 |
| N | 1.00 | 0.91 | 0.15 |
| MAC | 1.00 | 0.96 | 0.11 |
| MDC | 0.99 | 0.90 | 0.15 |
| Mono | 0.98 | 0.88 | 0.17 |
| MAST | 1.00 | 0.92 | 0.15 |
| cDC | 0.95 | 0.85 | 0.18 |
| pDC | 0.94 | 0.85 | 0.18 |
| cycMNP | 0.99 | 0.90 | 0.16 |
| cycNKC/T | 0.89 | 0.82 | 0.18 |

B

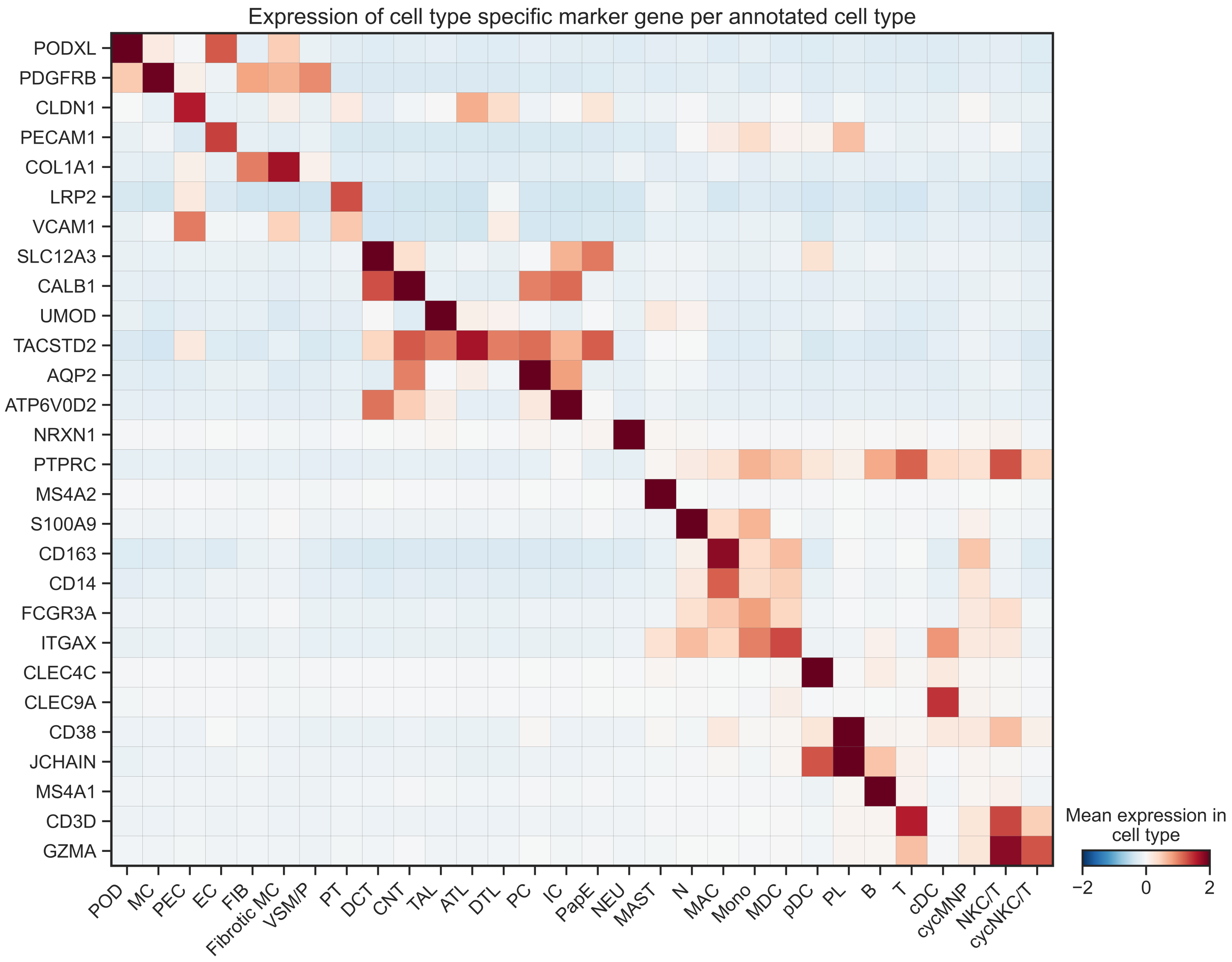

A

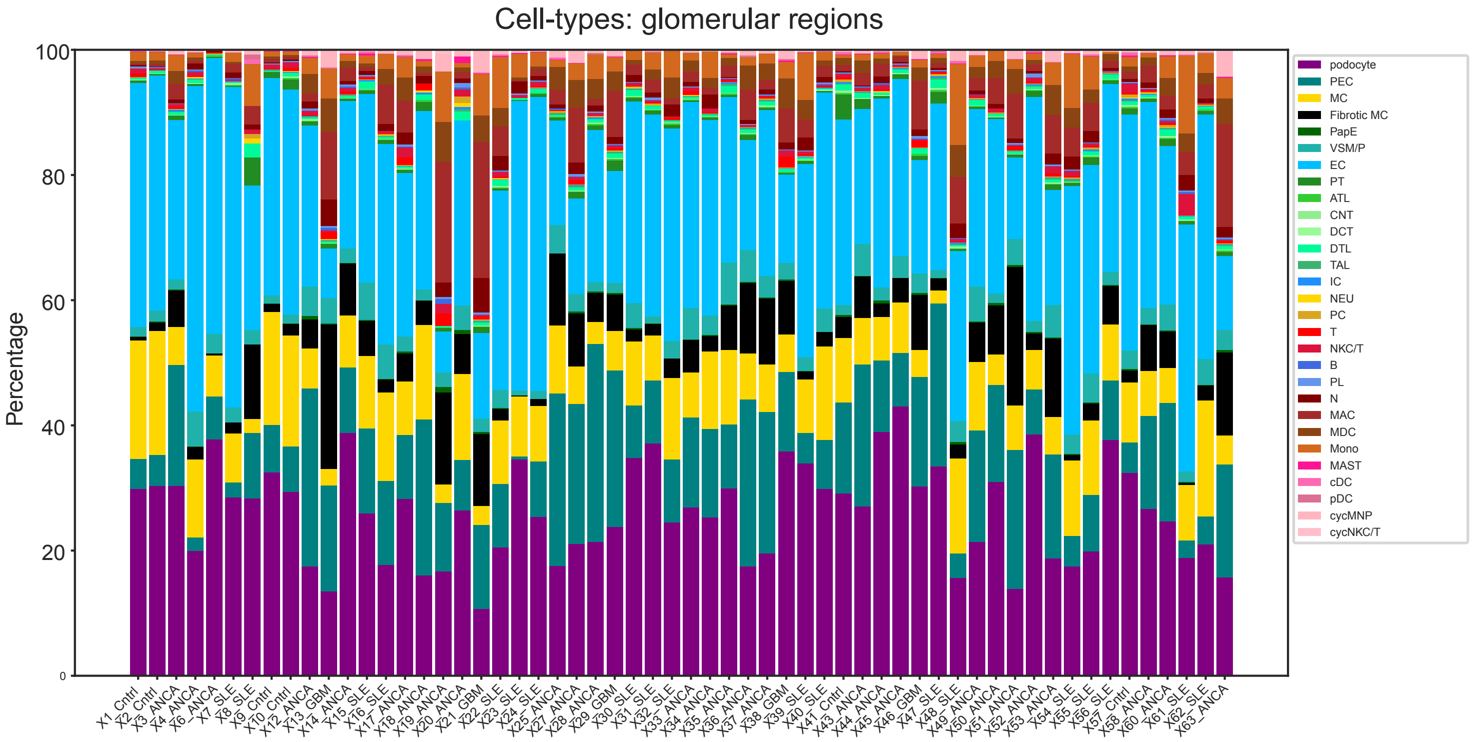
Cell-types: glomerular regions
Percentage
100
80
60
40
20
0
podocyte
PEC
MC
Fibrotic MC
PapE
VSM/P
EC
PT
ATL
CNT
DCT
DTL
TAL
IC
NEU
PC
T
NKC/T
B
PL
N
MAC
MDC
Mono
MAST
cDC
pDC
cycMNP
cycNKC/T
X1_Cntrl X2_Cntrl X3_ANCA X4_ANCA X6_ANCA X7_SLE X8_SLE X9_Cntrl X10_Cntrl X12_ANCA X13_GBM X14_ANCA X15_SLE X16_SLE X17_ANCA X18_ANCA X19_ANCA X20_ANCA X21_GBM X22_SLE X23_SLE X24_SLE X25_ANCA X27_ANCA X28_ANCA X29_GBM X30_SLE X31_SLE X32_SLE X33_ANCA X34_ANCA X35_ANCA X36_ANCA X37_ANCA X38_GBM X39_SLE X40_SLE X41_Cntrl X43_ANCA X44_ANCA X45_ANCA X46_GBM X47_SLE X48_SLE X49_ANCA X50_ANCA X51_ANCA X52_ANCA X53_ANCA X54_SLE X55_SLE X56_SLE X57_Cntrl X58_ANCA X60_ANCA X61_SLE X62_SLE X63_ANCA

B

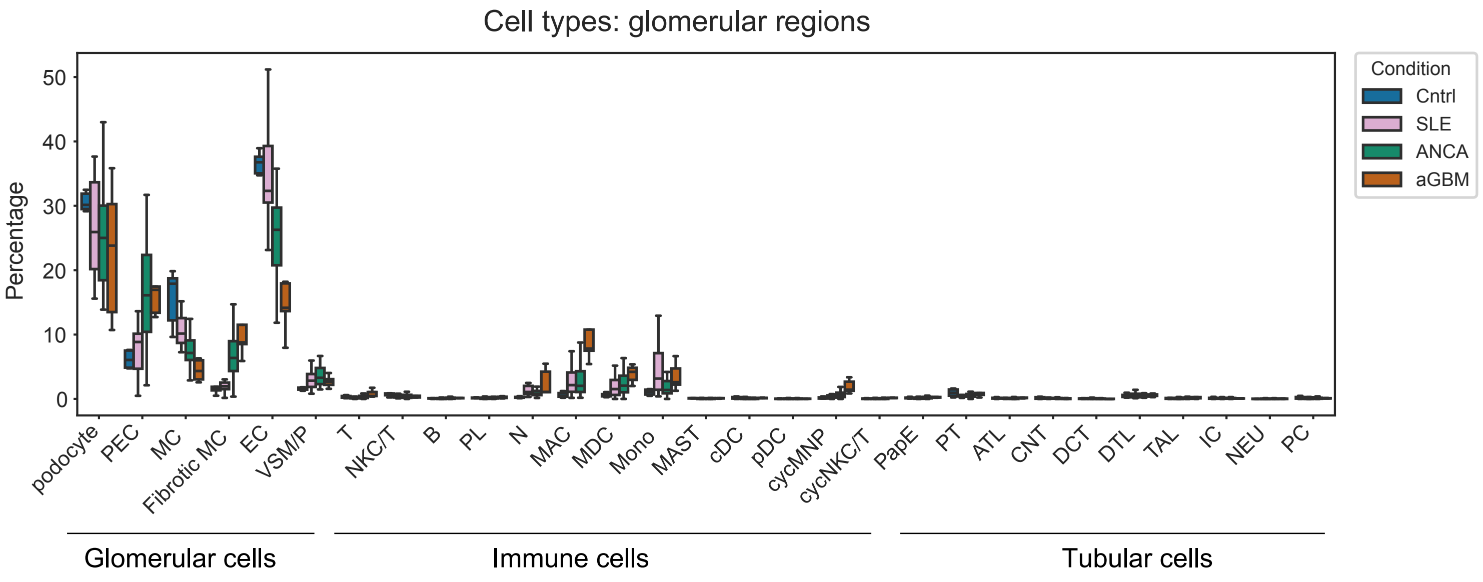
Cell types: glomerular regions
Percentage
50
40
30
20
10
0
Condition
Cntrl
SLE
ANCA
aGBM
podocyte PEC MC Fibrotic MC EC VSM/P T NKC/T B PL N MAC MDC Mono MAST cDC pDC cycMNP cycNKC/T PapE PT ATL CNT DCT DTL TAL IC NEU PC
Glomerular cells
Immune cells
Tubular cells

A

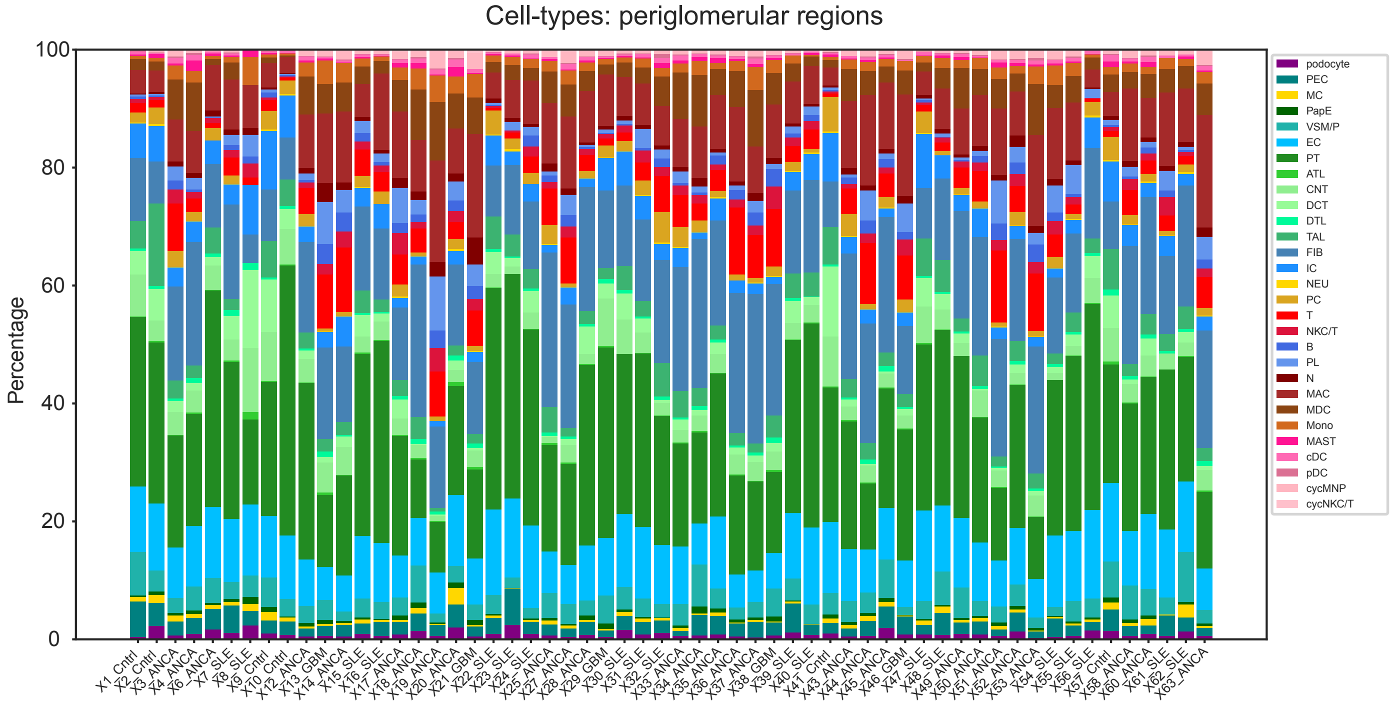
Cell-types: periglomerular regions
Percentage
0
20
40
60
80
100
podocyte
PEC
MC
PapE
VSM/P
EC
PT
ATL
CNT
DCT
DTL
TAL
FIB
IC
NEU
PC
T
NKC/T
B
PL
N
MAC
MDC
Mono
MAST
cDC
pDC
cycMNP
cycNKC/T

B

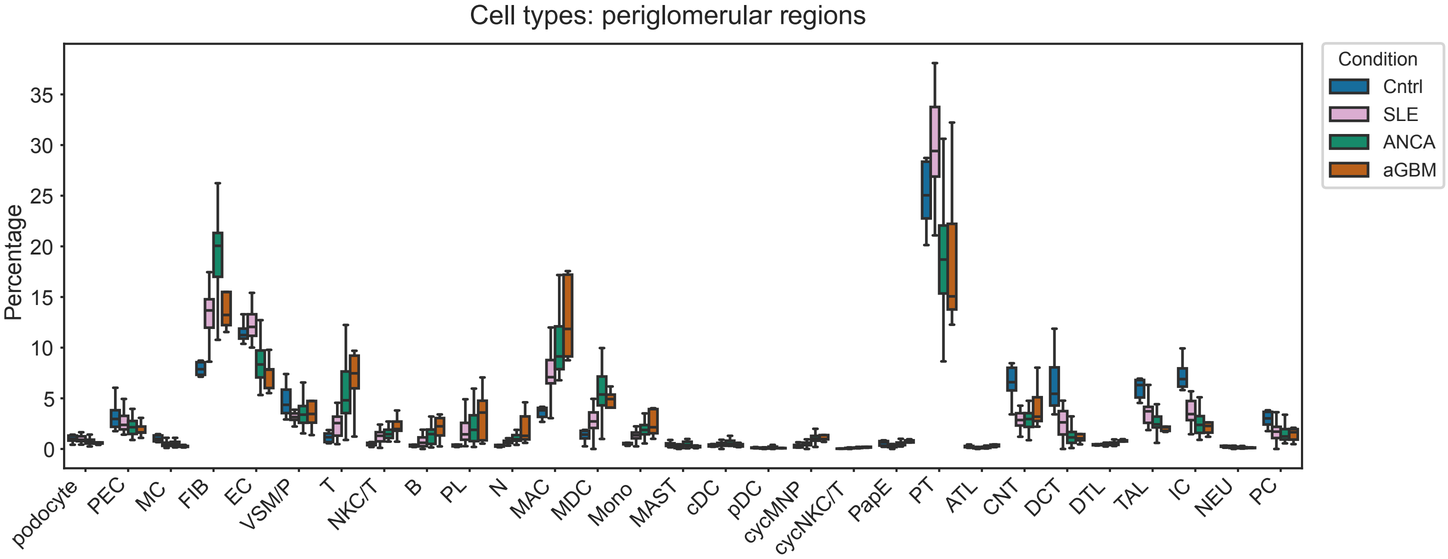
Cell types: periglomerular regions
Percentage
0
5
10
15
20
25
30
35
Condition
Cntrl
SLE
ANCA
aGBM
podocyte
PEC
MC
FIB
EC
VSM/P
T
NKC/T
B
PL
N
MAC
MDC
Mono
MAST
cDC
pDC
cycMNP
cycNKC/T
PapE
PT
ATL
CNT
DCT
DTL
TAL
IC
NEU
PC
Glomerular cells
Immune cells
Tubular cells

Extended Data Fig 5

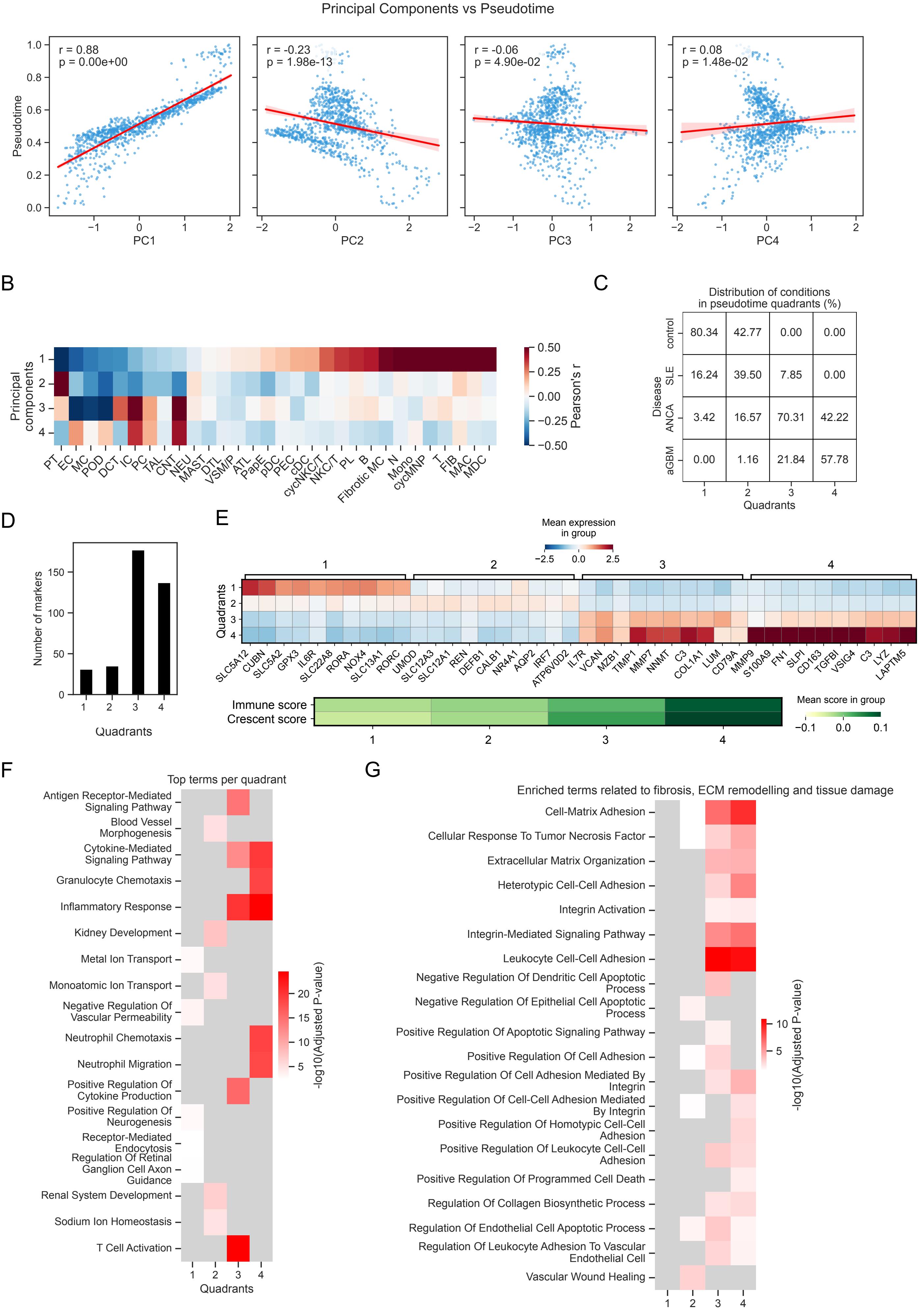

A

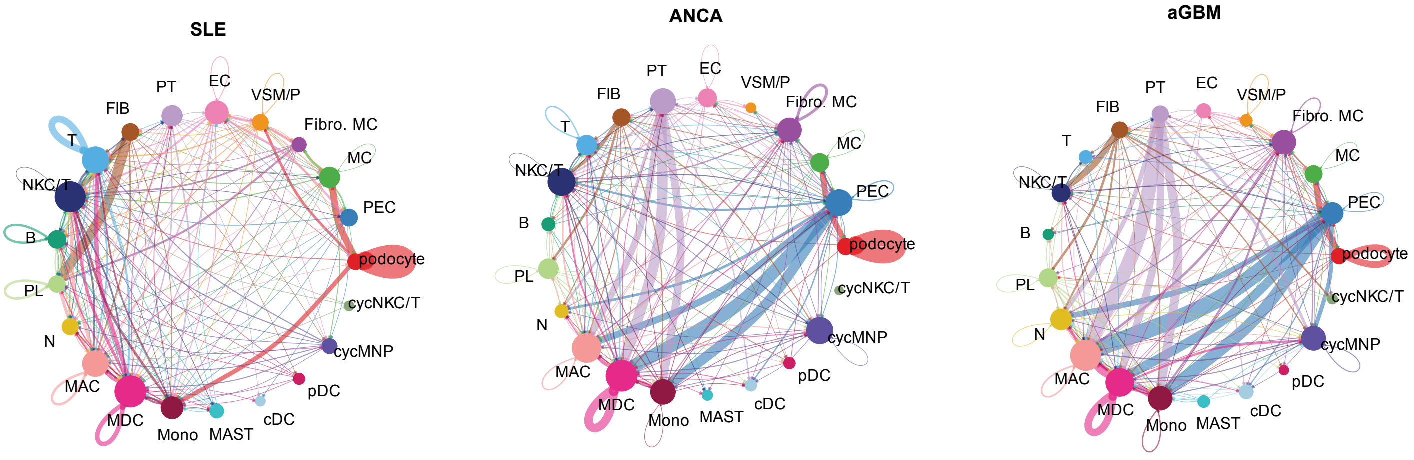
SLE
ANCA
aGBM
PT
EC
VSM/P
FIB
Fibro. MC
T
MC
NKC/T
PEC
B
podocyte
PL
cycNKC/T
N
cycMNP
MAC
pDC
MDC
cDC
Mono
MAST

B

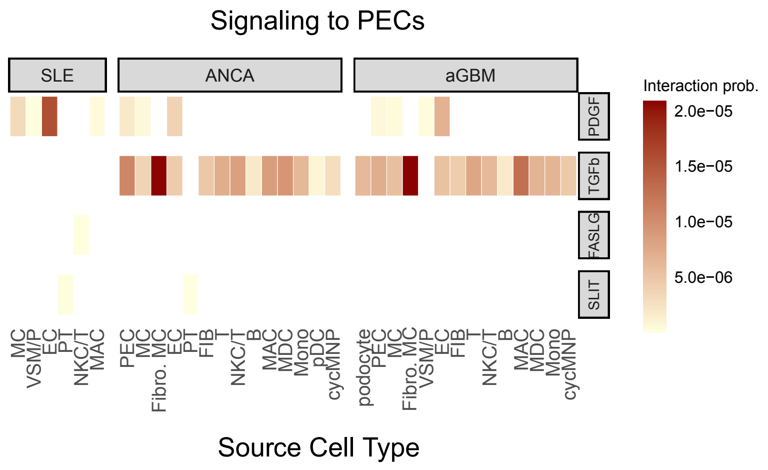
Signaling to PECs
SLE
ANCA
aGBM
PDGF
TGFb
FASLG
SLIT
Interaction prob.
2.0e−05
1.5e−05
1.0e−05
5.0e−06
MC
VSM/P
EC
PT
NKC/T
MAC
PEC
MC
Fibro. MC
EC
PT
FIB
T
NKC/T
B
MAC
MDC
Mono
pDC
cycMNP
podocyte
PEC
MC
Fibro. MC
VSM/P
EC
FIB
T
NKC/T
B
MAC
MDC
Mono
cycMNP
Source Cell Type

A

QC metrics: single nuclei RNA-sequencing

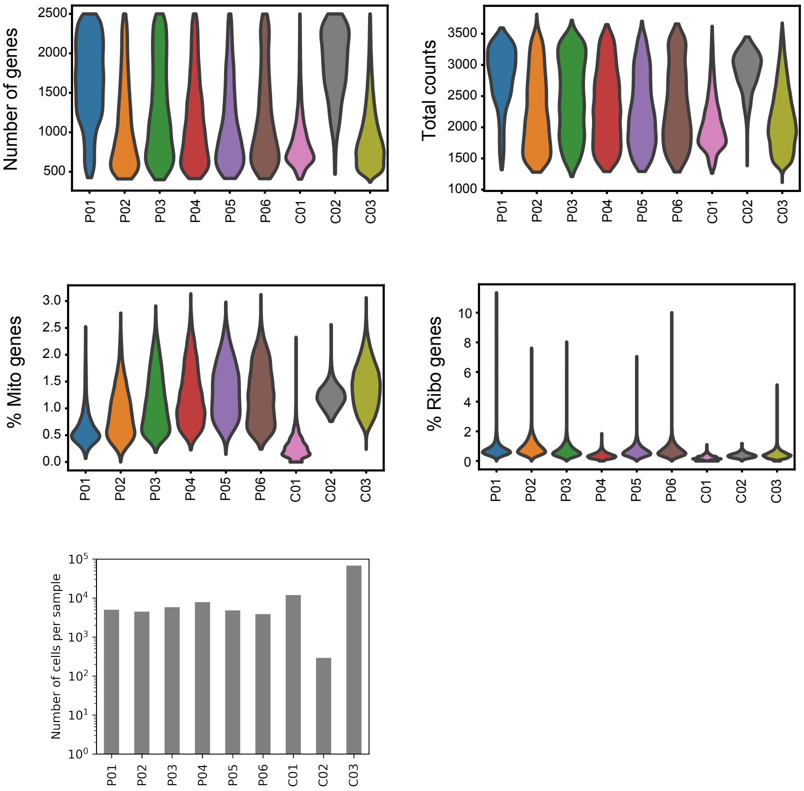


B

Cell type confidence

| | Median | Mean | SD |
|---|---|---|---|
| podocyte | 1.0 | 0.98 | 0.08 |
| mesangial cell | 1.0 | 0.94 | 0.13 |
| PEC | 1.0 | 0.96 | 0.11 |
| EC | 1.0 | 0.93 | 0.14 |
| FIB | 0.97 | 0.85 | 0.2 |
| VSM/P | 0.99 | 0.87 | 0.19 |
| PT | 1.0 | 0.94 | 0.13 |
| DCT | 1.0 | 0.9 | 0.17 |
| CNT | 0.95 | 0.85 | 0.18 |
| PC | 0.98 | 0.89 | 0.16 |
| IC | 1.0 | 0.94 | 0.13 |
| TAL | 1.0 | 0.94 | 0.13 |
| ATL | 0.64 | 0.68 | 0.22 |
| DTL | 0.93 | 0.84 | 0.18 |
| MAST | 0.53 | 0.55 | 0.19 |
| MAC | 0.95 | 0.86 | 0.18 |
| ncMON | 0.92 | 0.82 | 0.21 |
| MDC | 0.98 | 0.89 | 0.17 |
| PL | 0.52 | 0.55 | 0.18 |
| B | 0.89 | 0.77 | 0.3 |
| T | 1.0 | 0.94 | 0.13 |
| cycMNP | 1.0 | 0.93 | 0.14 |
| NKC/T | 0.92 | 0.86 | 0.16 |

C

Expression of cell type specific marker genes per cell type

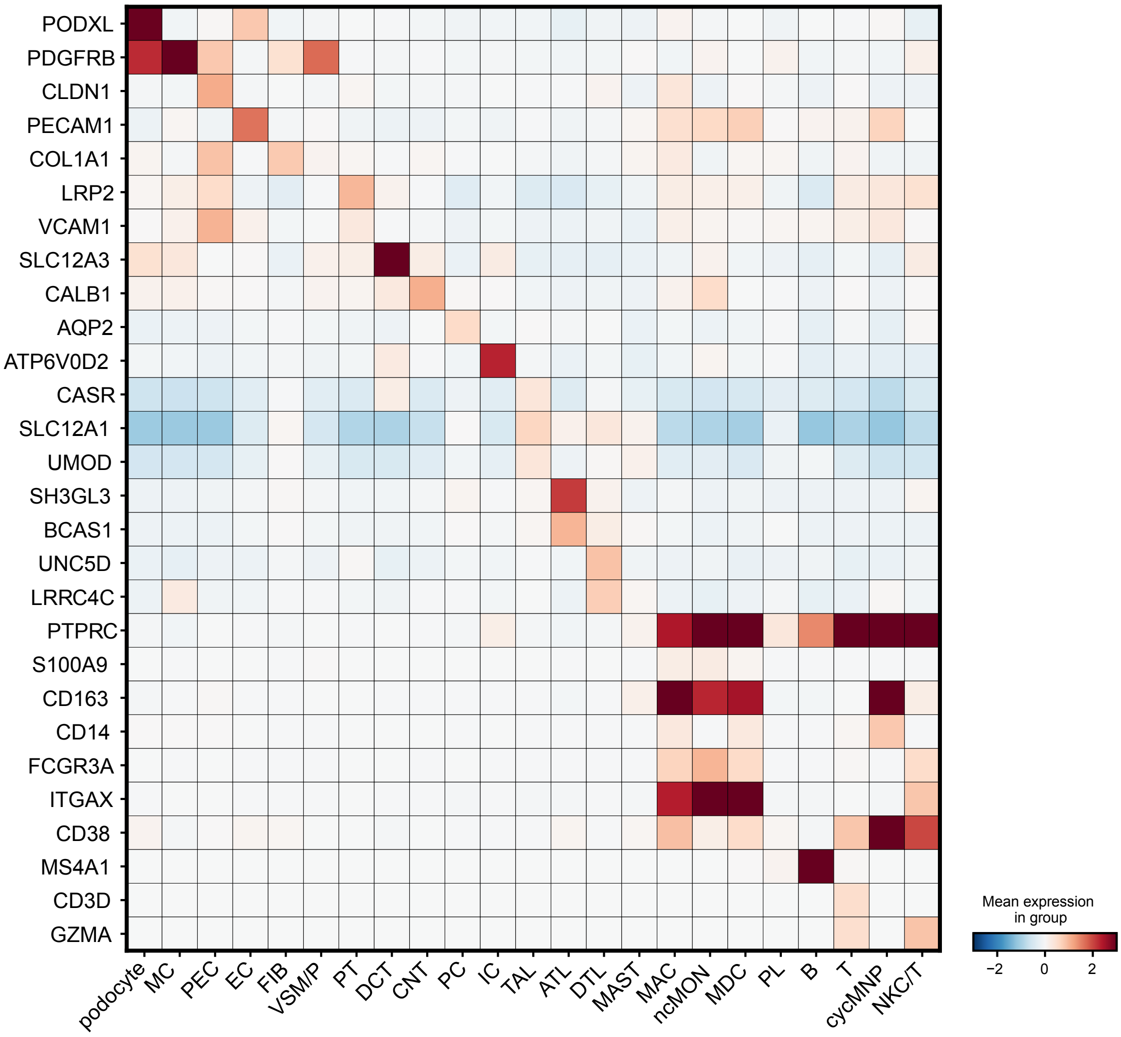

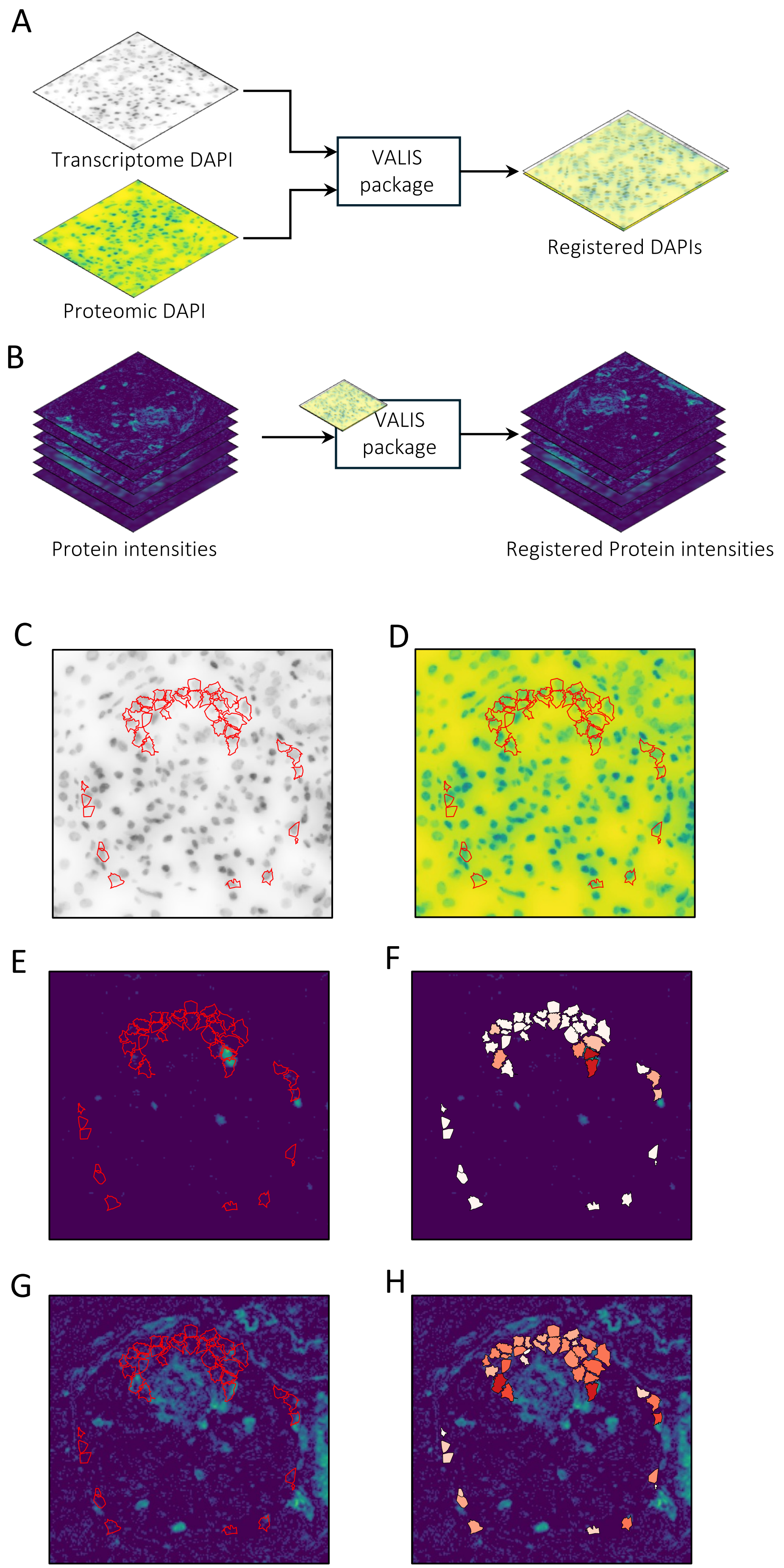
A
Transcriptome DAPI
Proteomic DAPI
VALIS package
Registered DAPIs
B
Protein intensities
VALIS package
Registered Protein intensities
C
D
E
F
G
H

### Extended Data Table 1: Gene Panel

| | | | | | | |
|---|---|---|---|---|---|---|
| ABCC3 | CD33 | CXCR6 | IL12RB1 | KLRD1 | PFKP | SLC8A1 |
| ABR | CD38 | CYBB | IL12RB2 | KLRF1 | PIGR | SLCO2A1 |
| ACKR3 | CD3D | DACH1 | IL13 | KLRG1 | PIK3CD | SLCO2B1 |
| ACSM3 | CD3E | DDIT3 | IL13RA1 | KLRG2 | PIK3CG | SLFN12L |
| ACTA2 | CD3G | DEFA1 | IL15 | KLRK1 | PLAUR | SLIT2 |
| ADAM17 | CD4 | DEFB1 | IL17A | LAG3 | PLCG2 | SLPI |
| ADAMTS5 | CD40LG | DEFB4A | IL17F | LAPTM5 | PLEC | SOD2 |
| AFDN | CD44 | DES | IL17RA | LATS2 | PLOD2 | SPI1 |
| AHR | CD69 | DNM1 | IL17RB | LCP2 | PLVAP | STAT1 |
| ALDH1A2 | CD74 | DOCK8 | IL17RC | LEF1 | PLXDC2 | STAT3 |
| ANGPT2 | CD79A | DPYSL3 | IL18 | LIF | PNP | STAT4 |
| APBB1IP | CD86 | DUSP6 | IL18BP | LRG1 | PODXL | STAT6 |
| AQP1 | CD8A | EDN1 | IL18R1 | LRP2 | POU2AF1 | STMN1 |
| AQP2 | CD93 | EGFL7 | IL1B | LSP1 | PPP1R15A | TACSTD2 |
| AREG | CD96 | EIF2AK1 | IL1R1 | LTB | PPP1R15B | TAGLN |
| ARG1 | CD99 | EIF2AK2 | IL1RL1 | LTBP2 | PRDM1 | TBX21 |
| ARHGAP15 | CDH13 | EIF2AK3 | IL2 | LUM | PRF1 | TCF7 |
| ASCL2 | CDH2 | EIF2AK4 | IL21 | LYVE1 | PRICKLE1 | TENT5C |
| ATF3 | CDH6 | EIF2S1 | IL21R | LYZ | PRKCH | TGFB1 |
| ATF4 | CDK4 | EMCN | IL22 | MAF | PROX1 | TGFBI |
| ATF5 | CDKN1A | ESAM | IL23A | MGP | PRTN3 | TGFBR1 |
| ATF6 | CEACAM1 | F11R | IL23R | MKI67 | PTGDR2 | TGFBR2 |
| ATP6V0D2 | CFH | FABP5 | IL24 | MMP1 | PTGS2 | THEMIS |
| BATF3 | CFTR | FAP | IL26 | MMP13 | PTPN22 | TIGIT |
| BCL11B | CHST11 | FAS | IL2RA | MMP14 | PTPN6 | TIMP1 |
| BCL6 | CIITA | FASLG | IL2RB | MMP3 | PTPRC | TKT |
| BHLHE40 | CLDN1 | FCER1A | IL2RG | MMP7 | PTPRD | TNC |
| BMP7 | CLEC10A | FCER1G | IL32 | MMP9 | RAC2 | TNF |
| BNC2 | CLEC4C | FCGR3A | IL33 | MPO | RAMP1 | TNFAIP2 |
| BTLA | CLEC9A | FCGR3B | IL3RA | MRC1 | RAMP3 | TNFRSF18 |
| C1QB | CLU | FCRL5 | IL4 | MS4A1 | RBCK1 | TNFSF10 |
| C1QC | CNN1 | FGF13 | IL4R | MS4A2 | REN | TNFSF11 |
| C3 | COL16A1 | FGFBP2 | IL5 | MSN | RGS1 | TNFSF13B |
| C4A | COL1A1 | FLT1 | IL5RA | MT2A | RIPK2 | TNFSF14 |
| C5 | COL3A1 | FN1 | IL6 | MUC5AC | RNASE2 | TNFSF8 |
| C5AR1 | COL6A3 | FOXN3 | IL6R | MUC5B | ROBO1 | TOP2A |
| C7 | COL7A1 | FOXP3 | IL6ST | MYO1G | RORA | TOX |
| CALB1 | COL8A1 | FPR3 | IL7R | MZB1 | RORC | TPSAB1 |
| CAMK1D | CORO1A | FUT4 | IL9 | NCAM1 | RUNX1 | TRAC |
| CCL18 | CPA3 | FUT7 | IL9R | NCR1 | RUNX3 | TRAF3IP2 |
| CCL19 | CREB5 | FYB1 | IRF4 | NCR2 | S100A6 | TRAF6 |
| CCL2 | CRK | GATA3 | IRF5 | NCR3 | S100A9 | TRAFD1 |
| CCL20 | CRKL | GNLY | IRF7 | NFATC2 | S1PR1 | TRAV1-2 |
| CCL21 | CSF1 | GPR183 | IRF8 | NKG7 | S1PR3 | TRAV24 |
| CCL26 | CSF1R | GPX3 | ITGA1 | NMI | S1PR4 | TRBC1 |
| CCL3 | CSF2 | GZMA | ITGA11 | NNMT | S1PR5 | TRBC2 |
| CCL4 | CSF2RA | GZMB | ITGA2 | NOD1 | SELE | TRDC |
| CCL5 | CSF2RB | GZMH | ITGA4 | NOS2 | SELENOK | TRGC1 |
| CCL7 | CSF3 | GZMK | ITGA8 | NOX4 | SELL | TRGC2 |
| CCN2 | CTHRC1 | HAVCR1 | ITGAE | NPHS2 | SELP | TYMP |
| CCR2 | CTLA4 | HAVCR2 | ITGAL | NPNT | SEMA4D | TYROBP |
| CCR3 | CUBN | HIF1A | ITGAM | NR4A1 | SEMA5A | UMOD |
| CCR4 | CX3CL1 | HMGB1 | ITGAX | NRG3 | SEMA6A | VASP |
| CCR6 | CX3CR1 | ICAM1 | ITGB1 | NRP1 | SERPINE1 | VCAM1 |
| CCR7 | CXCL1 | ICOS | ITGB2 | NRXN1 | SIGLEC8 | VCAN |
| CCR8 | CXCL10 | ICOSLG | ITGB8 | NT5E | SIPA1 | VEGFA |
| CCR9 | CXCL11 | ID3 | JAG1 | NTN4 | SIRPA | VIM |
| CD14 | CXCL12 | IFNA1 | JAM2 | OAS3 | SKAP1 | VSIG4 |
| CD163 | CXCL13 | IFNB1 | JAM3 | OLFML2B | SLC12A1 | WT1 |
| CD177 | CXCL16 | IFNG | JCHAIN | OXR1 | SLC12A2 | XCL1 |
| CD19 | CXCL2 | IFNGR1 | JUN | P2RX7 | SLC12A3 | XCL2 |
| CD1C | CXCL3 | IFNGR2 | KCNJ15 | PARP8 | SLC13A1 | XCR1 |
| CD2 | CXCL5 | IGFBP5 | KDR | PDCD1 | SLC14A1 | XG |
| CD200 | CXCL8 | IGFBP6 | KIRREL1 | PDE1C | SLC22A8 | ZAP70 |
| CD209 | CXCL9 | IKZF2 | KIT | PDGFA | SLC26A4 | ZEB2 |
| CD226 | CXCR2 | IL10 | KLF2 | PDGFB | SLC4A1 | ZNF683 |
| CD247 | CXCR3 | IL10RA | KLF3 | PDGFRB | SLC4A9 | |
| CD27 | CXCR4 | IL12A | KLRB1 | PDPN | SLC5A12 | |
| CD28 | CXCR5 | IL12B | KLRC1 | PECAM1 | SLC5A2 | |
